# Phosphorothioate DNA modification by BREX Type 4 systems in the human gut microbiome

**DOI:** 10.1101/2024.06.03.597175

**Authors:** Yifeng Yuan, Michael S. DeMott, Shane R. Byrne, Katia Flores, Mathilde Poyet, Mathieu Groussin, Global Microbiome Conservancy, Brittany Berdy, Laurie Comstock, Eric J. Alm, Peter C. Dedon

## Abstract

Among dozens of microbial DNA modifications regulating gene expression and host defense, phosphorothioation (PT) is the only known backbone modification, with sulfur inserted at a non-bridging oxygen by *dnd* and *ssp* gene families. Here we explored the distribution of PT genes in 13,663 human gut microbiome genomes, finding that 6.3% possessed *dnd* or *ssp* genes predominantly in Bacillota, Bacteroidota, and Pseudomonadota. This analysis uncovered several putative new PT synthesis systems, including Type 4 Bacteriophage Exclusion (BREX) *brx* genes, which were genetically validated in *Bacteroides salyersiae.* Mass spectrometric analysis of DNA from 226 gut microbiome isolates possessing *dnd*, *ssp*, and *brx* genes revealed 8 PT dinucleotide settings confirmed in 6 consensus sequences by PT-specific DNA sequencing. Genomic analysis showed PT enrichment in rRNA genes and depletion at gene boundaries. These results illustrate the power of the microbiome for discovering prokaryotic epigenetics and the widespread distribution of oxidation-sensitive PTs in gut microbes.

**One-sentence Summary:** Application of informatic, mass spectrometric, and sequencing-based mapping tools to human gut bacteria revealed new phosphorothioate epigenetic systems widespread in the gut microbiome.

## Introduction

Epigenetic modifications have been found in DNA from all domains of life. Among the variations in canonical A, T, C and G structure, such as *N*^6^-methyl-adenine (6mA), *N*^4^-methyl-cytosine (4mC), *C*^5^-methyl-cytosine (5mC), and 7-deazaguanine derivatives^1, 2^, phosphorothioates (PT) are the only known modification of the sugar-phosphate backbone, with a non-bridging oxygen replaced by sulfur in R_P_ specific configuration^3-5^ (**Fig. 1A**). Two PT-based restriction and modification (R-M) systems, the *dnd*^4, 6^ and *ssp*^7, 8^ gene clusters, have been observed in ∼10% of bacteria and archaea^9, 10^. Typically, the modification components *dndBCDE* and *sspBCD* are present in the form of a three- and four-gene operon, respectively. Like methylation-based R-M systems^8, 11, 12^, DndACDE and SspBCD proteins catalyze PT modification on one or both strands of specific consensus sequences. For example, DndACDE confer double-stranded PTs at 5′-G_PS_AAC-3′/5′-G_PS_TTC-3′ sequences in *Escherichia coli* B7A and *Salmonella enterica* serovar Cerro 87, 5ʹ-G_PS_GCC-3′/5′-G_PS_GCC-3′ in *Pseudomonas fluorescens* pf0-1 and *Streptomyces lividans* 1326, and 5′-G_PS_ATC-3′/5′-G_PS_ATC-3′ in *Hahella chejuensis* KCTC 2396^13-15^. SspABCD proteins, on the other hand, catalyze single-stranded 5′-C_PS_CA-3′ in *Vibrio cyclitrophicus* FF75^7, 13^. The *dnd* and *ssp* systems share some similarities, such as encoding a homolog of phosphoadenosine phosphosulphate (PAPS) reductase (DndC, SspD)^7, 16^, a homolog of cysteine desulfurase (DndA, SspA)^7, 16^, and a P-loop containing ATPase (DndD, SspC)^7, 17^ (**Fig. 1B**). The restriction counterparts DndFGH and SspFGH sense PT modifications by poorly understood mechanisms, with only 10-15% of consensus sequences modified with PTs^7, 13^.

**Figure 1.**
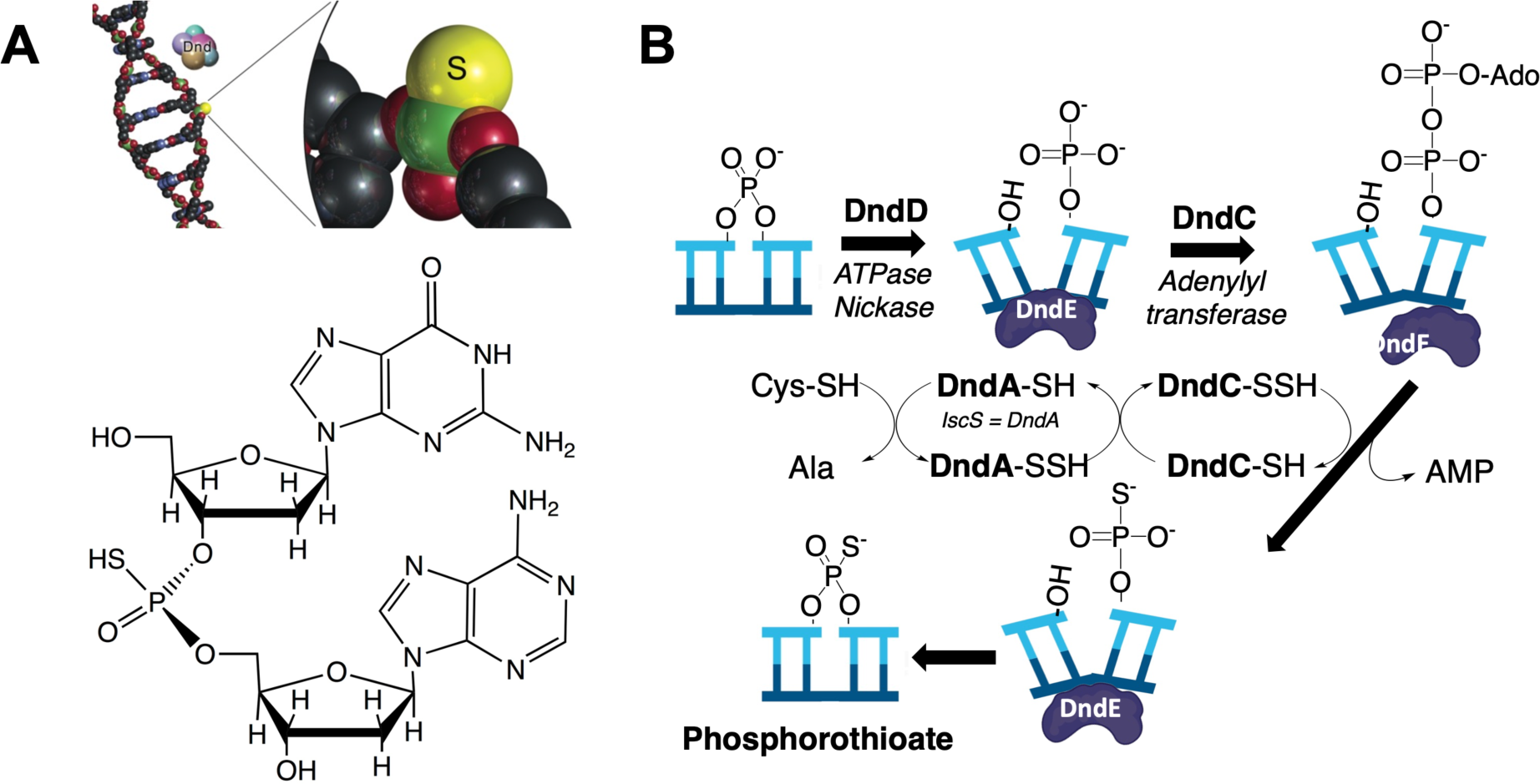
Microbial phosphorothioate (PT) DNA modifications. (**A**) Sulfur replaces a non-bridging phosphate oxygen in the DNA backbone in PT modifications. A limit nuclease digest of PT-containing DNA leaves PT-linked dinucleotides that can be identified and quantified by LC-MS. (**B**) Synthesis of PTs generally follows the biochemical steps performed by Dnd proteins.

Both the metabolic pathways enabling sulfur incorporation into PTs and the unique chemical properties of sulfur-modified DNA have implications for interactions of PT-containing microbes with human hosts. For example, the sulfur in PTs is both readily oxidized and nucleophilic, which is proposed to provide epigenetic regulation of transcription of redox homeostasis genes^15^. PTs also provide weak to modest protective effects in cells exposed to reactive oxygen and nitrogen species, such as peroxides ^15, 18^ and peroxynitrite^19^. Contrasting with this protection, PT-containing bacteria are 5-fold more sensitive to neutrophil-derived hypochlorous acid (HOCl) due to extensive DNA breaks at PTs ^20^. These unusual chemical properties of PTs raise questions about how PT-containing microbes might behave in the healthy gut microbiome or be altered by inflammatory bowel disease (IBD) or other chronic inflammatory conditions^21-23^. Evidence for the presence of PT genes in bacterial strains associated with the human gut microbiome and PT dinucleotides in fecal DNA^24, 25^ thus motivated us to systematically analyze PT genomics in the human gut microbiome.

Here we defined the landscape of PT-containing microbes in the human gut by performing a genomic analysis of 13,000 human microbiome genomes from the Broad Institute-OpenBiome Microbiome Library (BIO-ML)^26^, the Global Microbiome Conservancy (GMbC)^27^, and the Unified Human Gastrointestinal Genome (UHGG) collection^28^. Mass spectrometric analysis of PT dinucleotides in 226 of these isolates coupled with PT-specific next-generation sequencing (PT-seq)^29^ led to the discovery of a new PT system involving Type 4 Bacteriophage Exclusion (BREX) genes *brx PRXL*. These results expand our knowledge about the diversity of PT epigenetics and lay the foundations for understanding the role of PT-containing microbes in human health and disease.

## Results

### Discovery of new PT modification systems by analyzing genome neighborhoods of *dndC* and *sspD* genes

Given the power of physical clustering analyses to identify gene functions in bacteria, we first performed a comprehensive gene neighborhood analysis^30^ of *dnd* and *ssp* genes to find new PT modification systems. Here we used Enzyme Function Initiative (EFI) tools^30^ to first search Uniprot for DndC and SspD homologs in sequence similarity networks (SSN) followed by the EFI Genome Neighborhood Tool to identify the genomic contexts of the SSNs. DndC and SspD possess a PAPS reductase domain with ATP pyrophosphatase activity essential for PT biosynthesis (**Fig. 1B**), an activity shared by sulfur-inserting RNA modification enzymes ThiI^31^, MnmA^32^, and Ncs6^32^, and TtcA^33^. As a second criterion for PT synthesis, we required that DndC and SspD neighborhoods also possess a P-loop-containing NTPase gene. Both DndD and SspC possess P-loop-containing ATPase activity essential for PT synthesis (**Fig. 1B**). Using this approach, we retrieved 3,120 DndC homologs and 2,132 SspD homologs from Uniprot by BLAST (e-value cutoff 10^-5^) and searched genome neighborhoods encoding both a PAPS reductase domain and a P-loop NTPase (**Supplementary Tables S1, S2**).

The resulting gene neighborhoods revealed several novel putative PT-modifying gene candidates. Each black circle or node in **Figure 2** depicts a neighborhood containing *dndC* (Fig. 1A) or *sspC* (Fig. 1B), with clustering of nodes based on SSNs. Here it is clear that most of the *dndC-*containing gene neighborhoods fall within the largest cluster and possess additional *dnd* genes consistent with canonical Dnd-based PT synthesis (**Fig. 2A**, nodes with red outlines). However, several of the smaller clusters contained neighborhoods with a *dndC* gene, a P-loop NTPase, and other known defense genes, suggesting PT-based RM systems. In one instance, *dndC* and the NTPase gene lie near genes for a Bacteriophage Exclusion (BREX) type 4 system^34^ (**Fig. 2A**, yellow nodes). Five of six BREX systems possess a BrxC/PglY ATPase, a PglX DNA methyltransferase, and PglZ phosphatase. In BREX Type 4 clusters containing *dndC*, the PglX adenine methyltransferase is replaced with the DndC S-inserting PAPS reductase domain protein^34^. For example, in *Methanothermobacter sp.* (**Fig. 2A** yellow node, black outline), *dndC* is adjacent to an NTPase gene, a *pglZ* phosphatase gene, and a Lon-like protease domain-containing *brxL* gene typically found in BREX Type 1 and 4 systems. In *Fischerella thermalis* (**Fig. 2A**, yellow node, red outline), the full set of *dndBCDE* genes cluster with *brxL*, *brxZ (pglZ)* phosphatase, *sspB* nickase, and a methyltransferase. The synthesis of PTs in the DndC-containing BREX type 4 system was subsequently validated genetically and by LC-MS, as discussed shortly.

**Figure 2.**
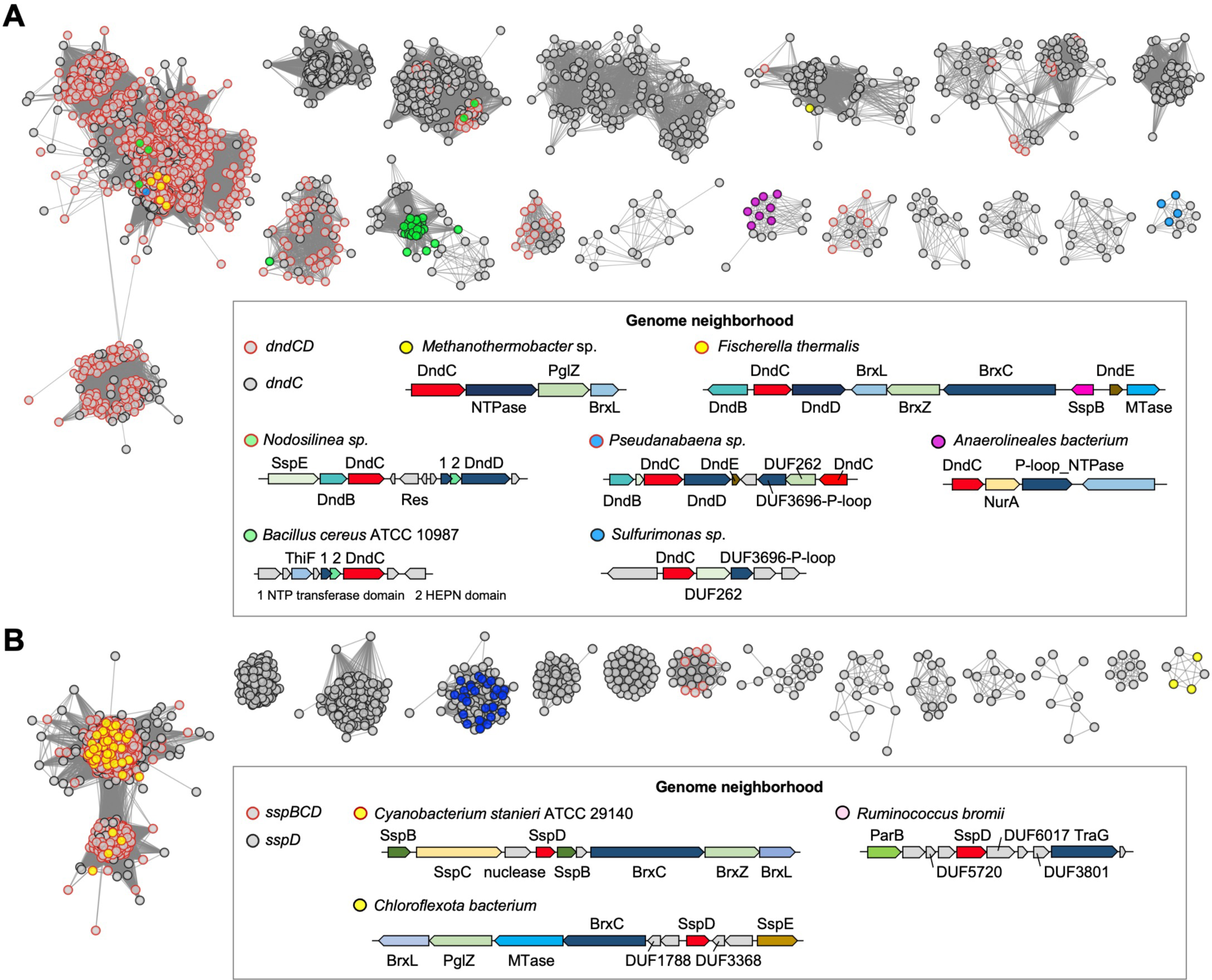
Gene neighborhoods analyses based on sequence similarity networks (SSNs) of DndC and SspD proteins essential for PT synthesis. SSN analysis was performed for the 3120 closest homologues of DndC (A) and 2132 homologues of SspD (B) in the UniProt database. Each node (black circle) in the network represents one DndC or SspD proteins. An edge (lines connecting nodes) is drawn between two nodes with a BLAST *E*-value cutoff of ≥10^–100^ (alignment score of 100) in the DndC SSN or 10^– 60^ (alignment score of 60) in the SspD SSN. The node outlines are colored according to the presence of other *dnd* or *ssp* genes, with red outlines denoting colocation of *dndC* with *dndD* and similarly for *sspD* with *sspBC*. Some nodes are colored according to the type of genome neighborhood structure, which are represented below for species discussed in the text. For better visualization, single nodes and clusters with a few nodes were hidden. Abbreviations: NTPase, Nucleoside triphosphatase; MTase, methyltransferase; Res, restriction enzyme; HEPN, higher eukaryotes and prokaryotes nucleotide-binding.

In a second putative PT defense system, a small cluster including *Bacillus cereus* (**Fig. 2A**, green node, black outline) pairs *dndC* with a putative minimal nucleotidyltransferase (MNT) and a higher eukaryotes and prokaryotes nucleotide-binding (HEPN) protein. This gene neighborhood resembles a Class II MNT-HEPN toxin-antitoxin (TA) system^35^, in which the HEPN protein is a RNase toxin, but its activity is neutralized by adenylylation by the MNT antitoxin^36, 37^. The HEPNs in the *dndC* clusters lack the RNase domain, but the neighboring MNT, ThiF, has the conserved motif GSX_10_DXD of an adenylyl transferase. Coupled with the adjacent NTPase, the DndC encoded in this Class II MNT-HEPN neighborhood could confer a three-component PT-based stress response system.

Finally, the *dndC* genes clustered with P-loop protein genes in the genome neighborhoods in two small clusters such as *Sulfurimonas sp.* (**Fig. 2A**, blue node, black outline) and *Anaerolineales bacterium* (**Fig. 2A**, pink node, black outline). A DUF262 gene was found in the fomer genome neighnorhood that is found in the SspE restriction component of Ssp-mediated PT systems and is involved in the PT-sensing anti-phage activity^7^.

The *sspD* gene neighborhoods were less complicated than those involving *dndC* (**Fig. 2B**). Again, *sspD* genes were associated with BREX defense system genes, as in the small cluster containing *Chloroflexota bacterium* (**Fig. 2B**, yellow node). This reinforces the idea that BREX type 4 genes represent a PT-based defense system.

To compare the distribution of *brx* genes to *dnd* and *ssp* families in prokaryotes, we performed a BLASTp search with protein sequences for DndABCDE, SspABCDE and BrxPCZL as queries in 6,616 representative genomes from the Bacterial and Viral Bioinformatics Resource Center (BV-BRC; as of January 2021)^38^. Here we defined the minimal genes necessary for a functional PT synthesis system as *dndCD*, *sspBCD*, *brxPCZL* based on the following observations: (1) DndA/SspA are often replaced by cysteine desulfurase such as IscS; (2) DndB is a non-essential regulator; (3) DndE protein is too short to search rigorously; (4) SspE is not required for PT modification; and (5) *brxPCZL* are the core 4 genes that define a BREX Type 4 system, with the *brxR* regulator not essential. Based on gene locus information for each protein hit, we found the *dndCD*, *sspBCD*, and *brxPCZL* gene clusters present in 4.3%, 3.0%, and 0.6%, respectively, of the BV-BRC genomes (**Supplementary Table S3**). Notably, both *dndCD* and *sspBCD* gene clusters co-occurred in Cyanobacteria, such as *Gloeocapsa* sp., *Coleofasciculus chthonoplastes*, and *Scytonema hofmanni*, among others (**Supplementary Table S3**). This complements the gene neighborhood analysis, which revealed that *dndC* clusters containing *brx* genes, MNT-HEPN genes, and DUF3696 genes are located near *dndBCD* operons in Cyanobacteria such as *Fischerella, Nodosilinea,* and *Pseudanabaena* (**Fig. 2A**, red-outlined yellow, green, and blue nodes in the main cluster). Similarly, 7 of 40 genomes with *sspBCD* operons located near *brxCZL* genes occurred in Cyanobacteria (**Fig. 2B**, red-outlined yellow nodes in the main cluster). These observations complement the observations of Lin *et al*.^9^ and suggest the co-evolution of the variety of PT-based epigenetic systems in the ancient photosynthetic Cyanobacteriota phylum.^39, 40^

### BREX Type 4 systems are homologous to Ssp proteins and catalyze PT synthesis

The genome neighborhood analyses revealed strong associations between the PAPS domain and NTPase genes essential for PT synthesis and *brx* genes from the BREX Type 4 family. This association was strengthened by the homology analysis (HHpred^41^) of BREX family and SSP proteins shown in **Figure 3A**, which reiterates the hallmark *brxPCZL* genes in BREX Type 4 and the lack of the PT-defining PAPS domain and NTPase genes in BREX Type 1. For example, BrxP possesses both the PAPS reductase domain found in SspD and the DUF4007 domain found in SspB (**Fig. 3A**), which effectively renders BrxP as a SspD-SspB fusion protein. Similarly, BrxC is essentially an SspC homolog, both possessing the same DUF6079 domain (**Fig. 3A**).

**Figure 3.**
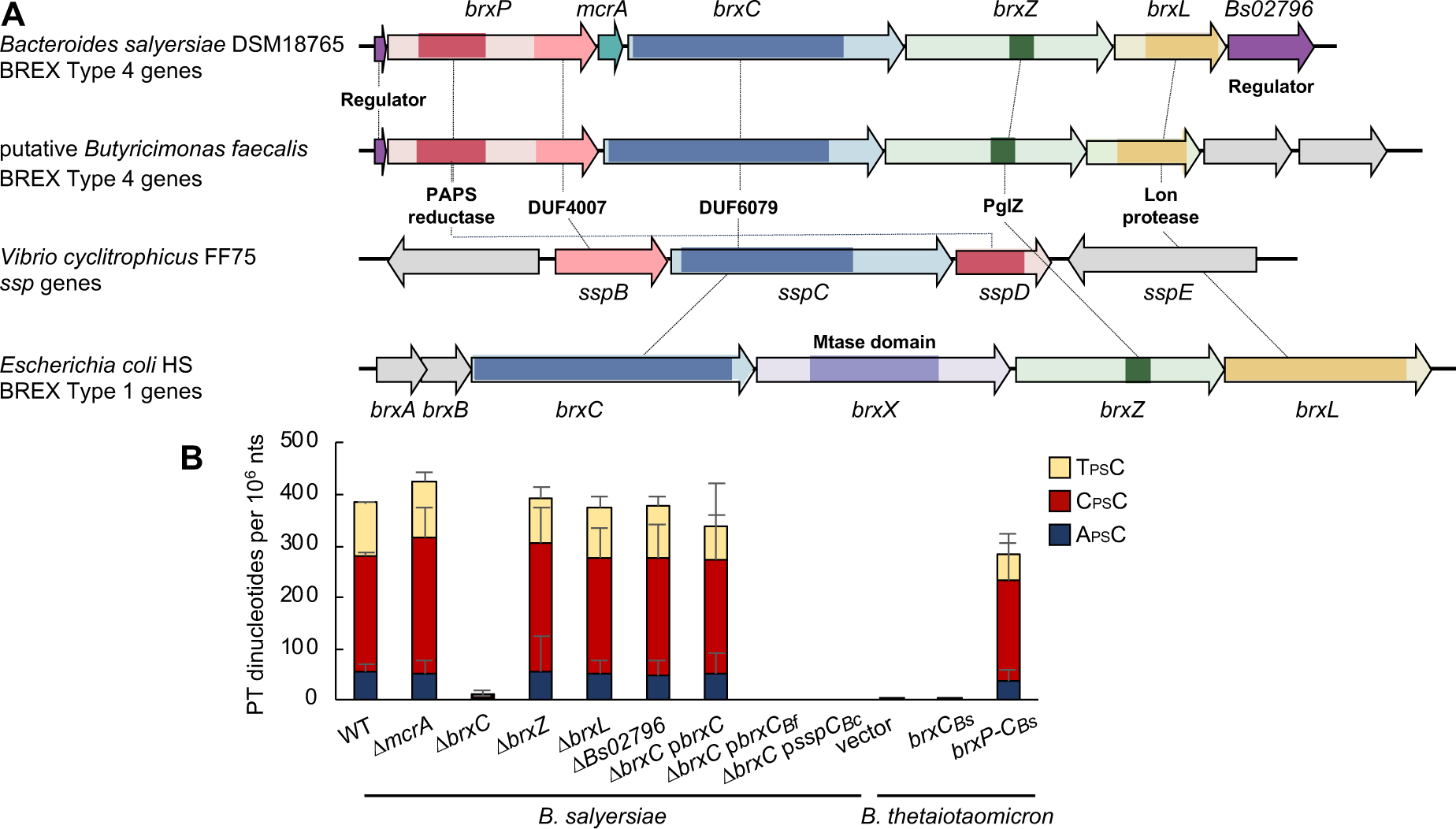
Homology analysis of BREX type 4 systems and evidence of PT synthesis. (**A**) The genomic organization of the BREX type 4 gene systems in *Bacteroides salyersiae* and putative *Butyricimonas faecalis*. *Vibrio cyclitrophicus* FF75 is included to demonstrate a typical *ssp* gene system and *Escherichia coli* HS to demonstrate a typical BREX type 1 system. Protein domains are color-coded and labeled. (**B**) The levels of PT dinucleotides in engineered *B. salyersiae* and *Bacteroides thetaiotaomicron* strains. *ΔbrxC, ΔbrxC* p*brxC_Bf_*, *ΔbrxC* p*sspCBc,* vector, and *brxCBs* are all significantly different from wilde-type (WT) by Student’s t-test, *p* <0.05.

Based on these similarities, we tested the PT synthesis activity of proteins encoded by genes in the BREX Type 4 family. The first step was to identify PT modifications in bacterial strains known to possess BREX Type 4 genes. Here we quantified PTs as PT-linked dinucleotides by liquid chromatography-coupled triple-quadrupole mass spectrometry (LC-MS/MS) in a limit digest with nuclease P1 and phosphatase dephosphorylation^42^. In *Bacteroides salyersiae* DSM18765, which harbors a 7-gene *brx* operon but no *dnd* or *ssp* genes (**Fig. 3A**), we detected the PT dinucleotides A_PS_C, C_PS_C, and T_PS_C at 56, 228, and 98 per 10^6^ nt, respectively (**Fig. 3B, Supplementary Fig. S1A, Supplementary Table S4**). As shown in **Supplementary Table S4**, several other bacterial isolates with a 4-5 gene complement of *brx* genes (*brxCLPRZ, brxCLPZ*) and insufficient sets of *dnd* or *ssp* genes also yielded C_PS_C (*Prevotalla sp*.), A_PS_A and A_PS_C (putative *Butyricimonas*), T_PS_C (*Parabacteroides*), and A_PS_A (*Bacteroides*). These results clearly distinguished the BREX Type 4 gene systems from *dnd* and *ssp* families.

Building on this associative evidence, we validated PT synthesis activity by creating in-frame deletion mutants of *brx* genes in *B. salyersiae*. As shown in **Figure 3B**, loss of *brxC* abolished PTs, with PTs restored by complementation with *in trans* expression of *brxC* from an expression plasmid. Based on the similarity between BrxP and SspD, we attempted to create a *brxP* deletion mutant with *B. salyersiae* but were unable to obtain this mutant, suggesting that its deletion may be deleterious. As an alternative, we reconstructed the PT pathway from *B. salyersiae* by cloning *brxP_Bs_, mcrA_Bs_* (a PT-dependent restriction enzyme in *Streptomyces coelicolor*)^43^, and *brxC_Bs_* or *brxC_Bs_* alone into the expression plasmid, which we transferred into *Bacteroides thetaiotaomicron* VPI, which lacks PT genes but possesses a SufS cysteine desulfurase homolog of DndA. The combination of *brxP_Bs_*, *mcrA_Bs_* and *brxC_Bs_* resulted in the T_PS_C, C_PS_C, and A_PS_C dinucleotides in the same proportion as wild-type *B. salyersiae.* While *brxC_Bs_* alone was not able to confer PTs, *mcrA_Bs_* was not required as shown in the *mcrA_Bs_* mutant (**Fig. 3B**). We conclude that, in presence of a cysteine desulfurase gene (*sufS*), *brxP* and *brxC* are the minimal set of BREX genes needed for PT synthesis.

### The distribution of PT modifying systems in the human gut microbiome

Based on the observation of PT-containing microbes in the mouse and human gut microbiome^24, 25^, we wondered about the presence of *brx*-based PT systems in gut bacteria and their relationship to *dnd* and *ssp* systems. To quantify the distribution of the PT systems in human gut microbiome, we searched for PT gene clusters in 13663 human gut microbial genomes from the BIO-ML^26^ and the GMbC^27^ collection, and from isolate sequences of human gut microbes and metagenome-assembled genomes (MAGs) in the Unified Human Gastrointestinal Genome (UHGG) collection^28^. As with the BV-BRC searches noted earlier, we performed a BLASTp search of proteins DndABCDE, SspABCDE and BrxPCZL as queries in these 13,663 genomes (**Supplementary Table S5**). The essential gene sets *dndCD*, *sspBCD*, *brxPCZL* were found to be present in 2.7%, 3.6%, and 1.4% of the humane gut microbiome genomes, respectively (**Fig. 4**). This represents a 1.6- and 1.2-fold reduction in *dnd* and *ssp* systems and a 2.3-fold enrichment in *brx* systems in the gut microbiome compared to the general BV-BRC genomes. The distributions of the three gene clusters at the genome level were nearly exclusive with few exceptions (**Supplementary Table S5**). Among the most prevalent phyla and orders in the human gut microbiome, the PT-modifying gene clusters were mainly found in Bacteroidota (Bacteroidales), Bacillota (Clostridiales), Pseudomonadota (Enterobacterales), and, to a lesser extent, Actinomycetota (Coriobacteriales) (**Fig. 4**). These observations raised the question of the predictive power of the genomic analyses.

**Figure 4.**
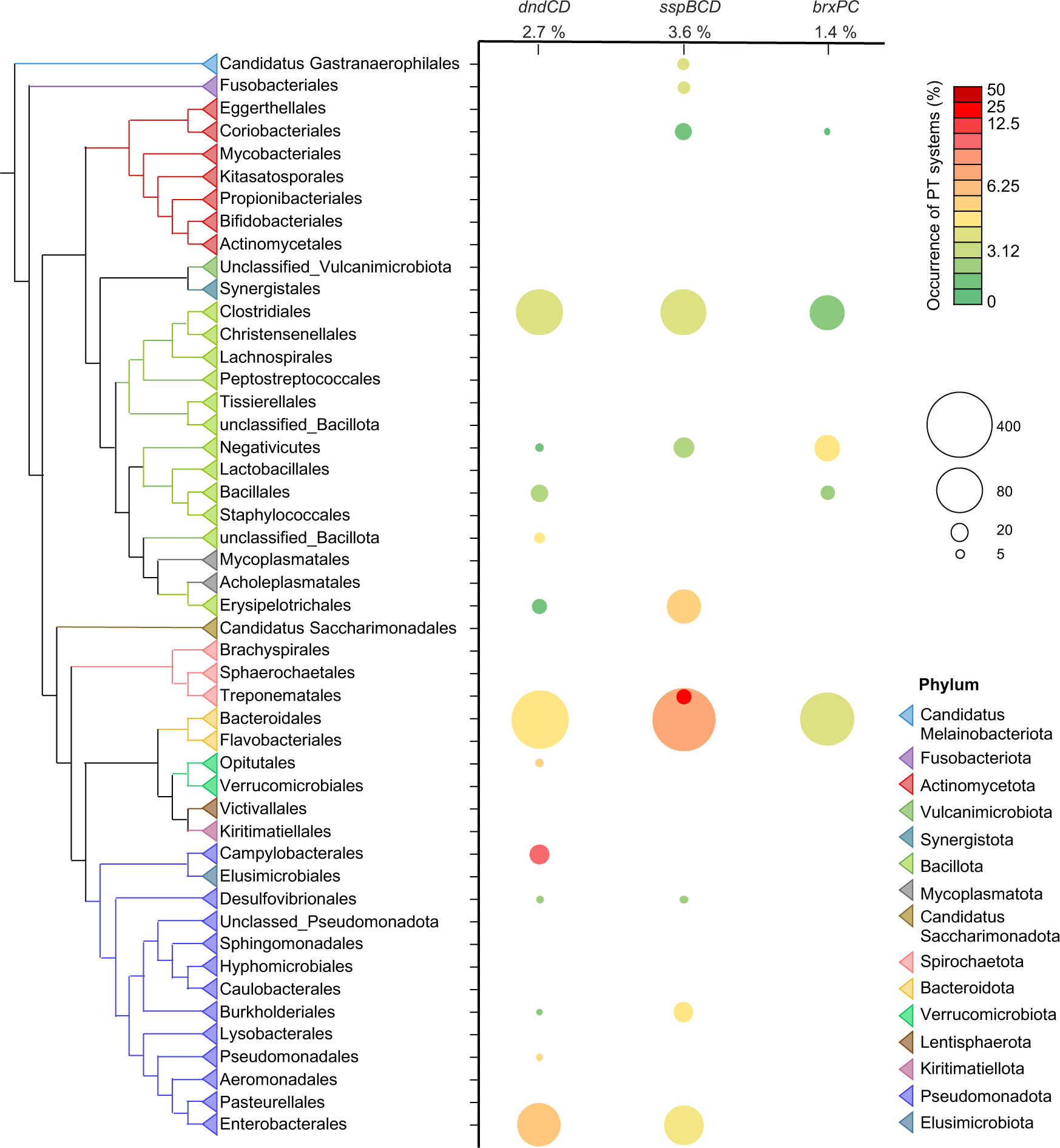
Phylogenetic distribution of *dndCD, sspBCD, and brxPC* genes in human gut microbiome genomes. The reference phylogeny was reconstructed from the concatenated alignment of 10 ribosomal proteins. The colors of the triangles in the tree show the taxa at the order level. The occurrence of gene clusters was quantitatively presented by the colors and size of circles. For better visualization, orders containing less than 5 genomes were hidden. Source data are provided as Table S5.

### Validating the predicted PT modifications in gut microbiome isolates

Given the presence of genes essential for PT synthesis in ∼1,000 out of 13,663 gut microbiome isolates, we next used LC-MS to validate the presence of PT dinucleotides in DNA in a collection of 226 bacterial isolates possessing *dndCD*, *sspBCD*, *brxPCZL* gene neighborhoods or individual PT synthesis genes not predicted to lead to PTs (**Supplementary Table S4**). In total, we identified 8 PT dinucleotides, including A_PS_A, A_PS_C, C_PS_A, C_PS_C, G_PS_A, G_PS_C, G_PS_T, and T_PS_C, and did not detect an additional two dinucleotides (C_PS_T, G_PS_G) that we found in human fecal DNA samples (**Supplementary Table S4**)^25^. Previous studies in three bacterial species with *dnd* and *ssp* genes^4, 42, 44^ revealed G_PS_A and G_PS_T at >500 per 10^6^ nt and C_PS_C at >2000 per 10^6^ nt, with C_PS_A, A_PS_A, A_PS_C, and T_PS_C detected much lower levels of 1-6 per 10^6^ nt^4, 42, 44^. The minor PT dinucleotides represent low-affinity binding sites for the DNA shape-selective Dnd proteins^44^. In sharp contrast, however, these minor sites in gut microbiome bacteria represent major PT modification sites, as high as 1,800 per 10^6^ nt in bacteria with *brxPCZL* genes (**Supplementary Table S4**).

The A_PS_A, A_PS_C, C_PS_A, C_PS_C, G_PS_A, G_PS_C, G_PS_T, and T_PS_C dinucleotides were distributed unevenly among the isolates based on the type of PT modification system, with four groups emerging from the analyses (**Fig. 4**). The first group of 39 isolates was characterized by G_PS_A and a predominance of *dnd* genes. The largest portion of Group 1 (32 isolates) possessed both G_PS_A and G_PS_T and harbored *dndCD* genes with or without *dndBE*, including isolates from Bacteroidales and Clostridiales orders. Five isolates from both Bacteroidales and Clostridiales harboring the minimal *dndCD* gene set showed G_PS_A and G_PS_C. One isolate that possessed a solitary *brxP* but no *dnd* genes and possessed both G_PS_A and G_PS_T. Given the presence of different dinucleotides for the *ssp* and *brx* gene families, this latter observation could be explained by genome sequencing errors or a sample mix up during regrowth, either resolved by re-sequencing. A single *Blautia* species harboring *dnd* genes showed only G_PS_A (**Supplementary Table S4**).

The second group of 34 isolates was characterized by the C_PS_C dinucleotide and *ssp* genes, with all but 2 harboring *sspBCD* ±*sspE* (**Fig. 5**). One isolate harbored *dndCD* genes that are not proximal to the *sspBCD* operon and showed LC-MS evidence of low levels of G_PS_A and G_PS_T at the detection limit of ∼2 PTs per 10^6^ nt (**Supplementary Fig. S2**), though these signals await high-resolution mass spectrometric validation. One isolate in this group lacks *sspBC* genes and one lacks all essential *sspBCD* genes. Again, the consistency of dinucleotide distributions suggests that *sspBCD* or *brxPCZL* genes are present in the genomes and perhaps missed due to insufficient genome sequencing coverage.

**Figure 5.**
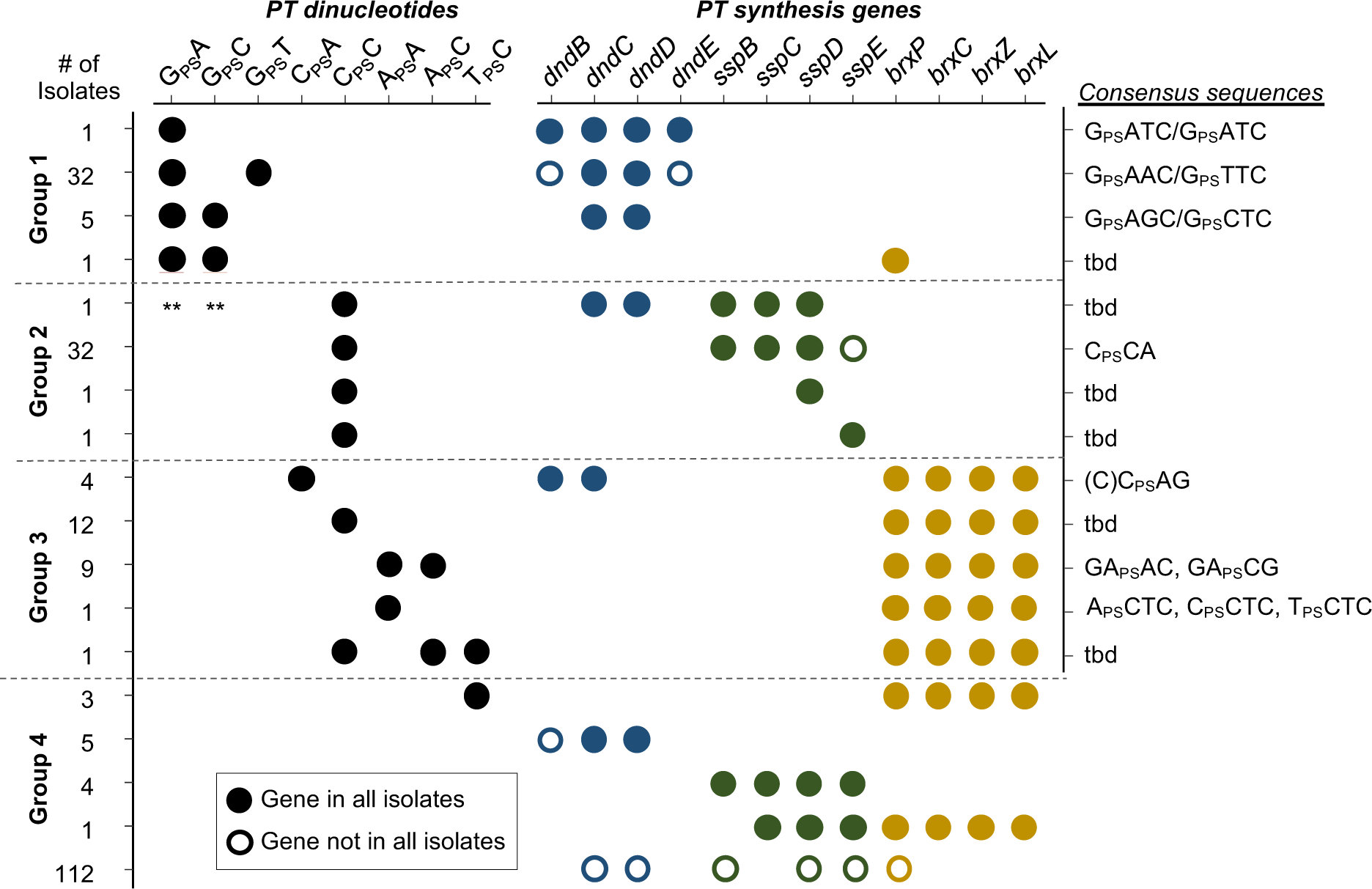
PT dinucleotides and PT consensus sequences in human gut microbiome isolates. The numbers of isolates were analyzed are listed on left. The circles represent PT dinucleotides quantified by LC-MS (left) and the presence of corresponding genes (right). The PT-modified consensus motifs were characterized in one representative isolate from each group using PT-seq. t.b.d., to be determined. *, the C_PS_AG motif was characterized using metagenomics PT-seq in another on-going study. **, G_PS_A and G_PS_T were detected using LC-MS QQQ but the exact mass was not verified by Orbitrap.

The third group of 30 isolates carried the BREX Type 4 genes *brxPCZL* and the most diverse sets of PT dinucleotides. Four isolates of *Parabacteroides* contain C_PS_A, 12 isolates of *Prevotella* sp. contain C_PS_C, 3 isolates of *Parabacteroides* contain T_PS_C, and 9 isolates of putative *Butyricimonas faecalis* contain A_PS_A and A_PS_C (Fig. S1B). The *B. salyersiae* strain examined earlier, which contains C_PS_C, T_PS_C, and A_PS_C (**Fig. 5, Supplementary Table S4**).

The fourth group of 122 bacteria lacked reliable LC-MS signals for PT dinucleotides. The majority (83) lacked the minimal sets of *dndCD*, *sspBCD* or *brxPC* gene clusters. For example, *Bacteroides dorei* CL03T12C01 harbors a PAPS reductase-encoding gene next to an ATPase without an adjacent *dndD* (**Supplementary Fig. S3**). Several isolates harbor *dndD* and *dptH* next to a methylase and a restriction enzyme but lack *dndC* (**Supplementary Fig. S3**). The absence of PT dinucleotides in these isolates agrees with the requirement for a PAPS reductase domain-containing protein and an NTPase. However, several (10) carry the minimal set of *dndCD* or *sspBCD* genes (**Fig. 4**), so PT dinucleotides were expected. Again, we cannot rule out errors in the reference genomes due to genome sequencing and sample handling.

These observations lead to a general conclusion that (1) Dnd proteins produce G_PS_A, G_PS_C, and G_PS_T, (2) Ssp proteins produce C_PS_C, and (3) Brx proteins produce everything except G_PS_A, G_PS_C, and G_PS_T. Understanding the basis for these differences requires knowledge about the longer consensus sequences containing the PT dinucleotides in the gut microbes.

### Novel PT consensus sequences in gut microbiome isolates

Here we defined novel larger PT consensus sequences, especially for the BREX Type 4 system, in gut microbiome isolates using a new and highly sensitive NGS sequencing technique, PT-seq^29^. The idea here is that the PT dinucleotides such as G_PS_A and G_PS_T, are found in the larger consensus sequences of G_PS_ATC and G_PS_AAC-3ʹ/5ʹ-G_PS_TTC in several types of bacteria^5^. Application of PT-seq (**Supplementary Fig. S4**) to the BREX Type 4-containing human gut microbiome isolate *B. salyersiae* DSM18765, earlier found to possess A_PS_C, C_PS_C, and T_PS_C at 62, 228, and 98 per 10^6^ nt (**Supplementary Table S4**), showed 5459 PT sites: 56 at A_PS_CTC, 3881 at C_PS_CTC, and 1513 at T_PS_CTC sites, respectively (**Fig. 6A**, **Supplementary Table S6**). As observed with other *dnd* and *ssp* systems, the Brx proteins modified only a portion of the 27274 ACTC (0.2%), 22698 CCTC (17%), and 36290 TCTC (4%) total sites available, respectively. The observation of PTs at 3 NCTC sites, with a strong preference for CCTC, is consistent with previous observations that Dnd proteins select their target sequences based on DNA shape rather than precise binding contacts^44^. In another BREX Type 4-containing putative *B. faecalis* (GMbC ID 5893AJ_0218_015_F2), PT-seq revealed 9832 GA_PS_AG and 5536 GA_PS_CG (**Supplementary Table S7**). These sites agreed with the A_PS_A and A_PS_C dinucleotides detected by LC-MS/MS (**Supplementary Table S4**). PT-seq also detected sites with bistranded PTs: 1080 GA_PS_AG/GT_PS_TG and 667 GA_PS_CG/CT_PS_GC. However, we could not detect reliable signals for the corresponding T_PS_T and T_PS_G dinucleotides by LC-MS/MS, which may be due to their low abundance and to insensitive MS detection of T-containing nucleotides.

**Figure 6.**
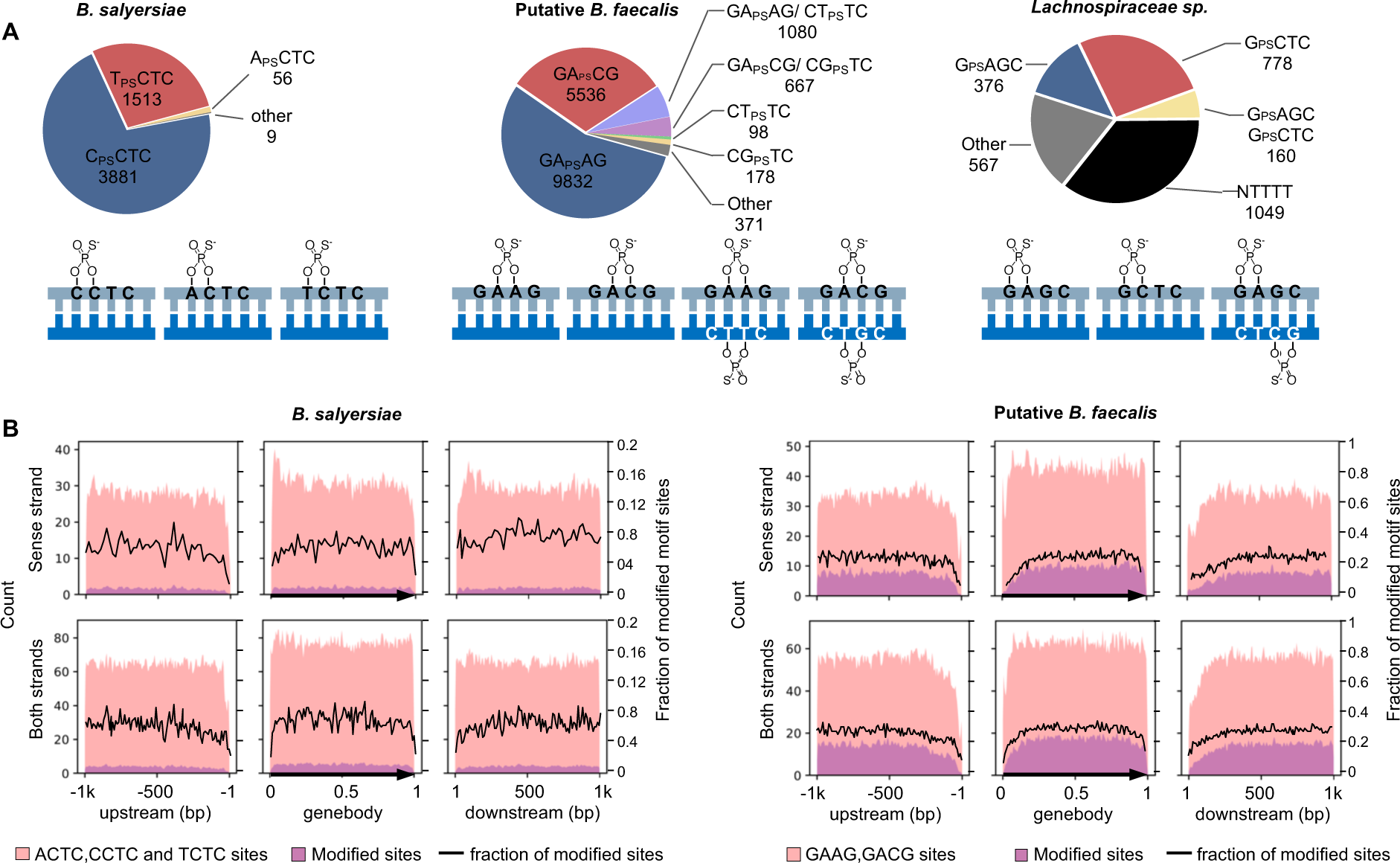
Biogeographical maps of PTs in bacteria with BREX type 4 systems. (**A**) Pie charts depict the number of single- or bi-stranded PT-modified consensus sequences. The structural diagrams depict these motifs. (**B**) Analysis of the distribution of PT consensus sequences and modified sites in 1kb upstream and downstream regions in *B. salyersiae* (**left**) and putative *B. faecalis* (**right**). The number of total motif sites in the sense strand (**upper**) or both strands (**lower**) are represented in pink. The PT modified sites are represented in purple. The fraction of PT-modified motif sites is represented by the black line.

Application of the original PT-seq protocol without biotin labeling and capturing to a *dnd* system-containing human gut microbiome isolate, *Lachnospiraceae sp.* (GMbC ID 2807EA_1118_063_H5), revealed 1,474 G_PS_AGC and G_PS_CTC sites occurring among the 16,683 possible GAGC/GCTC sites, with 1,154 modified on one strand and 160 modified on both strands (**Fig. 6A, Supplementary Table S8**). That a large fraction of pileup sites occurred at NTTT and other sites represent artifacts of poly(dT) tailing of single strand nicks using terminal transferase.

### The uneven genomic distribution of PTs

In addition to defining the consensus sequences for BREX Type 4 and other PT synthesis systems, PT-seq also revealed that PTs are nonrandomly distributed across bacterial genomes, which agrees with single-molecule real-time (SMRT) sequencing maps in *Escherichia coli*^13^. For example, the percentage of gene classes and intergenic regions containing PTs in *Lachnospiraceae sp., B. salyersiae, and B. faecalis* varied significantly: 6% of intergenic regions, 13% of tRNA genes, and 58% of coding sequences (CDS) (**Supplementary Table S9**). The percentage of consensus sequences modified with PTs, which is roughly 10% in genomes studied to date^13, 15^, also varied in individual bacterial genomes, ranging from 1% in tRNA genes in *Lachnospiraceae sp.* to 40% in rRNA genes in *B. faecalis* (**Supplementary Table S9**). Interestingly, the consensus sequences recognized by PT synthesis proteins were underrepresented at the boundaries of CDS (**Supplementary Fig. S5**). We further assessed the distribution of PTs within and between CDSs by quantifying PT-modified and unmodified consensus sequences in the sense strand of gene bodies and in regions 1 Kbp upstream and downstream of all CDSs. For both *B. faecalis* and *B. salyersiae,* this analysis revealed that the percentage of PT-modified consensus sequences was lower at the boundaries of coding regions (**Fig. 5B**, black line; **Supplementary Fig. S6**). Published PT-seq data for *E. coli* BW25113, which has a bistranded G_PS_AAC/G_PS_TTC PT consensus, also showed this biased distribution of PTs (**Supplementary Fig. S7A**). Thus, both the consensus sequences for PTs and the proportion modified with PT were underrepresented on either side of CDSs. While PTs have been proposed to interfere with transcription^13^, the percentage of PT-modified consensus sequences did not correlate with the level of transcription activity (**Supplementary Fig. S7B**), nor did the number of PTs in possible promoter regions (**Supplementary Fig. S7C**). The basis for these biased distributions remains to be defined.

## Discussion

Here we applied systems-level informatic, mass spectrometric, and sequencing tools to discover and characterize new PT modification systems in bacteria, with a focus on gut microbes. A rigorous neighborhood analysis of the key PT synthesis genes in *dnd* and *ssp* systems led to the discovery of several potential new PT modification systems, including MNT-HEPN, DUF262, and BREX Type 4 gene gene families. Genetic and bioanalytical validation of the BREX Type 4 system revealed the essentiality of *brxPC* genes for PT synthesis in bacteria with *brxPCZL* clusters. With the discovery of the *dnd* system in 2005^3^ and the *ssp* system in 2020^7^, the *brxPCZL* cluster represents the third established PT modification system.

The extent of distribution of the *dnd*, *ssp*, and *brx* gene clusters was assessed first among 6,616 representative prokaryotic genomes and then among 13,663 gut microbiome isolates. The difference in the frequency of *dndCD*, *sspBCD*, *brxPCZL* gene clusters in the BV-BRC general set of bacteria (4.3%, 3.0%, and 0.6%, respectively) and the gut microbiome isolates (2.7%, 3.6%, and 1.4%, respectively) suggests a bias for the BREX Type 4 systems in the gut environment. The total of 7.7% of gut microbiome isolates possessing PT-synthesis genes is consistent with the estimate of 5-10% of stool microbes possessing PT modifications, as we observed using mass spectrometric measurement of PT dinucleotides in human fecal DNA^25^.

With the discovery of BREX Type 4 as the third PT synthesis gene family, one feature of PT modifications appears to be universal for PT epigenetics: partial modification of available consensus sequences in individual genomes. In agreement with previous sequencing studies with bacteria containing Dnd proteins^13, 45^, PT-seq analysis revealed that the BREX Type 4 proteins also only partially modify their four-nucleotide consensus sequences, ranging from 0.2% at ACTC to 17% at CCTC in *B. salyersiae* (**Fig. 5A, Supplementary Table S6**). The BREX Type system also points to another emerging feature of PT modification systems: sequence-selectivity of the *dnd, ssp*, and *brx* gene families. Previous^7, 8, 46^ and present studies (**Fig. 4**) consistently show that *ssp* systems catalyze C_PS_C dinucleotides mainly in CCA motifs, while the *dnd* systems insert PTs mainly in GN motifs: G_PS_A, G_PS_C, G_PS_G^42^, and G_PS_T dinucleotides. The latter occur mainly as bistranded modifications of GNNC motifs. We previously showed that Dnd proteins select modification sites based on DNA shape, with GAAC, GTTC, and GATC all sharing similar shapes and all being modified with PTs at G_PS_A and G_PS_T in *S. enterica*^44^. The fact that *ssp* is selective for CCA suggests a similar shape selectivity. BREX Type 4-dependent PT systems are more complicated, which may reflect the hybrid nature of the PT-synthesizing gene families in different bacteria. The five types of PT nucleotides produced by Brx proteins (C_PS_A, C_PS_C, A_PS_C, A_PS_A, T_PS_C) share C_PS_C with *ssp* systems but lack the G_PS_N unique to *dnd* systems (**Fig. 5A**). This may be partly explained by the sequence similarities between SspBCD and BrxPC proteins. The shape selectivity argument is again supported with *brxPCZL* in *B. salyersiae* DSM18765, with the ACTC, CCTC, and TCTC sites in an NCTC motif all modified to differing extents (**Fig. 5A**) and in *B. faecalis* with GAAG and GACG motifs.

Our studies revealed the complexity of interactions among the three PT systems, with widespread combinations of the various components of all three systems and coexistence of two or more gene clusters in the same organism (**Fig. 1**, **Supplementary Table S3**). These evolutionary co-occurrences likely reflect a fitness advantage, such as that observed for the presence of Dnd and Ssp together, which provides complementary and synergistic protection against temperate and lytic phages as well as phage induction^46^. The co-occurrence of combinations of all three established PT systems and two putative systems in Cyanobacteria supports the hypothesis of an evolutionary divergence that may originate from a sulfur-based metabolism in ancient Cyanobacteria ancestor^9^. All three PT systems contain a gene encoding a PAPS reductase domain protein that has been proposed to involve in the initial sulfur mobilization step^5^. The observation of a biased toward *brx*-based PT systems and the unique biochemical properties of PTs raise questions about the role of PT epigenetics in the human gut microbiome in health and disease.

## Methods

### Bacterial strains and growth conditions

Bacteria revived from BIO-ML and GMbC were cultured as previously decribed^26^. *Parabacteroides* spp. and *Bacteroides* spp., *E. faecalis*, and BIO-ML isolates were grown on ASF plates (Becton Dickinson) and BHIS plates^47^, BHI plates (Becton Dickinson), and Brucella agar, (Catalog # AS-141, Anaerobe Systems, Morgan Hill, CA), respectively. Culture manipulations were performed at 37 °C in an anaerobic chamber with an atmosphere of 80% nitrogen, 5% carbon dioxide, and 5% hydrogen. Growth on agar plates was harvested with phosphate buffered saline (Catalog # 10010-023, Gibco, Paisley, PA). The bacterial suspensions were pelleted by centrifugation at 10,000 rpm for 15 min at ambient temperature. The supernatant was removed and the pellets were stored at -20 °C.

Bacterial stains and plasmids used and generated for BREX 4 system deletion and reconstruction are listed in **Supplementary Table S10**. Bacteroidales strains were grown in basal liquid medium^48^ or on BHIS plates^47^. Antibiotics used for selection include erythromycin (10 µg/mL), gentamicin (200 µg/mL), anhydrotetracycline (50 ng/mL). *E. coli* S17 λ pir was grown in LB broth or plates with carbenicillin (100 µg/mL) added for selection.

### Creation of deletion mutants and complementing clones

Internal non-polar deletion mutants of *brx* genes HMPREF1532_02792, HMPREF1532_02793, HMPREF1532_02794, HMPREF1532_02795, and HMPREF1532_02796 were constructed by amplifying DNA upstream and downstream of each gene using the primers listed in **Supplementary Table S11**. These flanking pieces were cloned into BamHI-digested pLBG13^47^ using NEBuilder (New England BioLabs) and transformed into *E. coli* S17 λ pir. PCR confirmed plasmids were sequenced (Plasmidsaurus) to confirm correct error-free PCR amplification. The correct construct was conjugally transferred from *E. coli* into *B. salyersiae* and cointegrates were selected on gentamycin/erythromycin. Double recombination cross-outs were selected on BHIS plates with anhydrotetracycline (aTC, 50 ng/ml) and screened via PCR for mutant genotype.

Genes expressed *in trans* in *B. thetaiotaomicron* and *B. salyersiae* Δ2793 were PCR amplified and cloned into BamHI-digested pFD340^49^ using NEBuilder. Transformants were PCR screened and the plasmid was sequenced. Plasmids were conjugally transferred from *E. coli* to *B. salyersaie* and transconjugants were selected on gentamycin/erythromycin plates.

### Sequence Similarity Networks

Sequence similarity networks (SSNs) were generated by submitting the sequences of *E.coli* B7A DndC (GeneBank AIF62362.1) and *V. cyclitrophicus* FF75 SspD (NCBI Refseq WP_016789110.1) to the EFI-EST webtool^30^ using the BLAST option (e-value cutoff 10^-5^, maximum number of sequences retrieved 5000). The initial SSN was generated with an alignment score cutoff set such that each connection (edge) represents a sequence identity of approximately 40%. Sequences that share 100% sequence identity were grouped into a single node. More stringent SSNs were created by increasing the alignment score cutoff in small increments (usually by 5–10). This process was continued until each cluster is estimated to be isofunctional, in which the nodes represent enzymes that catalyze the same reaction. Isofunctionality was determined by mapping the known *dnd* clusters and following their movements as the alignment cutoff increased until the *dnd* clusters fell into subclusters. The network were visualized in alignment score weighted Prefuse Force-Directed Layout using Cytoscape 3.9^50^. The resulted SSNs were submitted to EFI-EST webtool^30^ for genome neighborhood analysis with default setting. Nodes were highlighted in SSNs if the neighboring Pfam families, with maximal median distance of no more than 4 and minimal co-occurrence rate larger than 0.2, have NTPase activity, NTP binding, or DNA binding activities.

### Sequence analyses

For sequence analyses, the BLAST tools^51^ (with e-value cutoff of 10^-10^ and query coverage of 25%) and HHpred^41^, and the resources of BV-BRC ^38^ were routinely used. In total, the data from 20279 bacterial and archaeal genomes were retrieved from the BIO-ML^26^, GMbC^27^, Unified Human Gastrointestinal Genome (UHGG)^28^, and BV-BRC representative bacteria as collected in Jan 2021. The queries used in this study were listed in **Supplementary Table S12**, including IscS (GeneBank AIF64277.1), DndB (GeneBank AIF62361.1), DndC (GeneBank AIF62362.1), DndD (GeneBank AIF62363.1), DndE (GeneBank AIF62364.1) in *E.coli* B7A and SspA (Refseq WP_016789103.1), SspB (Refseq WP_022570853.1), SspC (Refseq WP_016789109.1), SspD (Refseq WP_016789110.1), SspE (Refseq WP_016789111.1), and BrxP (GMbC tag OIFBNFKG_02855), BrxC (GMbC tag OIFBNFKG_02856), BrxZ (GMbC tag OIFBNFKG_02857), BrxL (GMbC tag OIFBNFKG_02858). The genes *dndCD*, *sspBCD*, and *brxPCZL* were considered to be present when all genes were adjacent.

### Phylogeny tree

We first built a concatenated alignment of 10 nearly universal and single-copy ribosomal protein families. We used Diamond v0.8.22^52^ (with parameters blastx -more-sensitive -e 0.000001 -id 35 -query-cover 80) to BLAST all proteomes in out collection against the RiboDB database v1.4.1^53^ of bacterial ribosomal protein genes. We excluded proteins bL17, bS16, bS21, uL22, uS3 and uS4, as they were not sufficiently distributed across all genomes. In each RiboDB gene family, we excluded genomes that contained gene duplicates. Then, we aligned all protein families individually with MUSCLE v5.1^54^ (with default setting). We filtered out misaligned sites using BMGE v1.12^55^ (with parameters -t AA -g 0.95 -m BLOSUM30) and concatenated all individual alignments using Seaview v4.7^56^. The phylogenomic tree was reconstructed using FastTree v2.1.10^57^ (with parameters -lg -gamma) and visualized and modified in iTOL^58^.

### DNA isolation

The cells were resuspended in 500 μL of phosphate buffered saline upon receiving and re-pelleted by centrifugation at 6,000 g for 5 min at 4 °C. Genomic DNA was isolated using E.Z.N.A Bacterial DNA kit (Catalog # D3350-02), with the bead beating option (speed of 4 m/s, 45 s “on”, 1 min rest, 3 cycles). DNA was eluted with 150-200 μL of RNase free water and stored at -80 °C.

### Digestion of DNA for LC-MS/MS analysis of PT dinucleotides

DNA (20 μg, 78 μL) was incubated with Nuclease P1 (1.5 U, 3 μL, US Biological) in 30 mM ammonium acetate pH 5.3 and 0.5 mM ZnCl_2_ (90 μL total reaction volume) for 2 h at 55 °C. The reaction mixture was diluted with Tris-HCl (100 mM final concentration, pH 8.0, 9 μL) and incubated with calf intestinal alkaline phosphatase (51 U, 3 μL, Sigma) for 2 h at 37 °C. Enzymes were removed by passing the mixture through a VWR 10 kDa spin filter with centrifugation at 12,000 g for 12 min. The solution was lyophilized to dryness and resuspended in H_2_O (50 μL).

### LC-MS/MS analysis of DNA PT dinucleotides

Synthetic PT DNA dinucleotides or nuclease P1 hydrolyzed DNA were analyzed by LC-MS/MS on an Agilent 1290 series HPLC system equipped with a Synergi Fusion RP column (2.5 μm particle size, 100 Å pore size, 100 mm length, 2 mm inner diameter) and a DAD. The HPLC was coupled to an Agilent 6490 triple quadrupole mass spectrometer. The column was eluted at 0.35 mL/min at 35 °C with a linear gradient of 3-9% acetonitrile in 97% solvent A (5 mM ammonium acetate pH 5.3) over 15 min. The column was rinsed with 95% acetonitrile in solvent A for 1 min, and then the initial conditions were regenerated by rinsing the column with 97% solvent A for 3 min. Canonical deoxyribonucleosides that eluted from the column were quantified by their 260 nm absorbance with the DAD. PT-containing dinucleotides were identified and quantified by tandem quadrupole mass spectrometry with electrospray ionization operated with the following parameters: N2 temperature, 200 °C; N2 flow rate, 14 L/min; nebulizer pressure, 20 psi; capillary voltage, 1800 V; and fragmentor voltage, 380 V. For product identification, the mass spectrometer was operated in positive ion multiple reaction monitoring mode using the conditions tabulated in **Supplementary Table S13**.

### High-resolution mass spectrometry

Synthetic PT dinucleotides (2 pmol per 10 μL injection) or Nuclease P1 hydrolyzed RNA (4 μg per 10 μL injection) were analyzed on a Dionex Ultimate 3000 UHPLC system equipped with a Synergi Fusion RP column (2.5 μm particle size, 100 Å pore size, 100 mm length, 2 mm inner diameter). The HPLC was coupled to a Thermo Fisher Q Exactive Hybrid Quadrupole-Orbitrap mass spectrometer. The column was eluted at 0.35 mL/min at 35 °C with a linear gradient of 3-9% acetonitrile in 97% solvent A (5 mM ammonium acetate pH 5.3) over 15 min. The column was rinsed with 95% acetonitrile in solvent A for 1 min, and then the initial conditions were regenerated by rinsing the column with 97% solvent A for 3 min. High resolution mass spectra for the PT-containing dinucleotides were obtained by hybrid quadrupole-Orbitrap mass spectrometry with the following parameters: sheath gas flow rate, 50 L/min; aux gas flow rate, 15 L/min; sweep gas flow rate, 3 L/min; spray voltage, 4.20 kV; and capillary temperature, 275 °C. For product identification, the mass spectrometer was operated in positive ion targeted single ion monitoring mode using the conditions tabulated in **Supplementary Table S14**.

### PT-seq library preparation

PT-seq applied on *Lachnospiraceae sp.* was adapted from previously described version^45^ (**Supplementary Fig. S4B**). 10 ug of genomic DNA was diluted in 500 μl of ddH2O in 2ml Eppie tube on ice and fragmented by probe sonication for 5 cycles (2 min ON with 50% duty cycle and ∼20 output control and 1 min OFF). The samples were subjected to SpeedVac concentration for ∼2.5 h at ∼450 mTorr to the final volume of ∼40 μL. Blocking of pre-existing strand-break sites was achieved by 4 cycles of denaturation-dephosphorylation-blocking. Each cycle started by denaturating at 94 °C for 2 min and immediately cooling down on ice for 2 min. The initial dephosphorylation reaction was in a mixture (50 μL) containing 5 μl of terminal transferase buffer (NEB Tdt Reaction Buffer, Catalog # M0315S), 1 μL of shrimp alkaline phosphatase (rSAP, NEB Catalog # M0371S), and 5 μg of fragmented DNA, with incubation at 37°C for 30min to remove phosphate at 3′ end of the strand-breaks. The phosphatase was then inactivated by heating at 65 °C for 10 min. After cooling, 1 μL of Tdt Reaction Buffer, 6 μL of CoCl_2_ (0.25 mM), 2 μL of ddNTPs (2 mM each, TriLink) and 1 μL of terminal transferase (20 units, NEB Catalog # M0315S) was added to the reaction with incubation at 37 °C for 1 h to block any pre-existing strand-break sites. Blocking cycles were repeated 3 times, in which fresh reagents were added in each cycle. The blocked DNA was purified using a DNA cleanup kit (Zymo Catalog # 11-304C).

For iodine cleavage, 32 μL of the blocked DNA was incubated with 4 μL of 500 mM Tris-HCl pH 9.0 and 4 μL of iodine solution (50 mM, Fluka, Catalog # 318981-100) at room temperature for 5 min. Then, the reaction product was purified using two DyeEX columns (QIAGEN Catalog # 63206) to remove salts and iodine. The purified DNA was denatured by incubating at 94 °C for 2 min and cooling on ice for 2 min and then incubated with 5 μL of NEB rCutsmart buffer and 1 μl of rSAP (50 μL reaction) to remove 3′-phosphates arising from iodine cleavage. After incubation at 37°C for 30 min and 65 °C for another 10 min, the product was incubated with 1 μL of dTTP (1 mM, NEB Catalog # N)443S), 1 μL of Tdt buffer, 6 μL of CoCl_2_, 1 μL of Tdt and 1 μL of H_2_O. After incubation at 37 °C for 45 min, the dTTP was removed by DyeEX columns.

PT-seq applied on *B. faecalis and B. salyersiae* was adapted by adding biotin labeling and streptavidin bead-capturing (**Supplementary Fig. S4A**), to resolve the inadequate poly(dT) tailing of iodine cleaved nicks. DNA (10 μg) was subjected to blocking of pre-existing strand-break sites, iodine cleavage, and dTTP labeling as described above. DNA (60 μL) was then incubated with 7.8 uL of Tdt buffer, 7.8 uL of CoCl_2_, 1 μL of ddUTP-biotin (1 mM, Jena Bioscience NU-1619-BIOX-S), 1 μL of Tdt and 1 μL of H_2_O at 37 °C for 1 h to terminate the T-tails by ddUTP-biotin. After cleaned with DyeEX columns, the DNA was diluted in 500 μL of H_2_O and fragmented by probe sonication as described above. The DNA fragments was mixed with 10 μL of streptavbidin coated meganetic beads (NEB Catalog # S1421S) and 500 μL of binding buffer (5 mM Tris-HCl, pH7.5, 1 M NaCl, 0.5 mM EDTA) and incubated on a shaker at ambient temperature for 1 h. The beads were pull down by a magnetic and washed 3 times with 100 μL of binding buffer. After discarding the supernatant, the beads were resuspended in 20 μL of H_2_O.

The purified product and captured beads in upgraded ICDS method were subjected to Illumina library preparation with SMART ChIp-seq kit (Takara, Catalog # 634865) by following the manufacturer’s protocol. The final step of PCR was performed using the Illumina primers provided in ChIp-seq kit and 12 cycles were used for amplification. The PCR product with unique sequencing barcode was submitted to Illumina MiSeq and NovaSeq instrument for 150 bp paired-end sequencing.

### Data analysis

As illustrated in the workflow (**Fig. S4**), the data analysis started with triming of Adapters. Adapters were removed using bbduk from BBtools (sourceforge.net/projects/bbmap/), (with parameters ktrim=r k=18 hdist=2 hdist2=1 rcomp=f mink=8 qtrim=r trimq=30 for R1, ktrim=r k=18 mink=8 hdist=1 rcomp=f qtrim=r trimq=30 for R2). T-tails were removed using bbduk with parameters ktrim=r k=15 hdist=1 rcomp=f mink=8 for R1 and ktrim=l k=15 hdist=1 rcomp=f mink=8 for R2. PhiX were removed using bbduk with parameters k=31 hdist=1. Trimmed reads were aligned to the corresponding genome using Bowtie 2 with setting “sensitive”. The bam files were cleaned using SMARTcleaner^59^ and split into two strands using samtools^60^. The coverage of each strand was calculated separately using bedtools^61^ with the genomecov -d -5 option. Then, custom shell scripts were used for pileup calling. Briefly, all read start positions were recorded for both strands separately. The read pileup depth at each position was represented by the number of read starting (5ʹ-end) at each position. The 13 nt sequences centered at positions with depth ≥1 were retrieved using samtools^60^. The consensus motifs were analyzed with incrementing depth, typically from 1 to 100 with step of 10, using MEME^62^ with parameters -dna -objfun classic -nmotifs 5 - mod zoops -evt 0.05 -minw 3 -maxw 6 -markov_order 0 -nostatus -oc. The read starting sites mapping within less than 3 bps were collapsed into the centremost consensus motif sites. Regions where starting positions mapped within 4 bp on opposite strands and within reverse complementary sequences were considered as double stranded modifications.

## Data availability

Custom scripts for processing the sequencing data are described in Methods and are available upon request. Sequencing data have been deposited in NCBI SRA database under BioProject ID PRJNA1006039.

## Acknowledgements

We thank Susan Weir and Katya Moniz at the Openbiome for providing us with bacterial isolates. This work was supported by National Institutes of Health (R01 ES031576), by a NIEHS Training Grant in Environmental Toxicology T32-ES007020 (S.R.B), and by funding for the GMbC from the MIT Center for Microbiome Therapeutics and the Neil and Anna Rasmussen Family Foundation.

## Notes

https://www.ncbi.nlm.nih.gov/sra

### Competing Interest Statement

The authors have declared no competing interest.

