## Supplementary Information: Figures and Tables for "Phosphorothioate DNA modification by BREX Type 4 systems in the human gut microbiome"

**Supplementary Information for**
**“Human gut microbiome comparative genomics reveals new phosphorothioate epigenetics systems”**

Yifeng Yuan, Michael S. DeMott, Shane R. Byrne, Katia Flores, Mathieu Groussin, Mathilde Poyet, Brittany Berdy, Laurie Comstock, Eric J. Alm, and Peter C. Dedon

***Contents***

• **Supplementary Figure S1**. Mass spectrometric validation of PT dinucleotides.
• **Supplementary Figure S2.** LC-MS/MS QQQ characterization of PT dinucleotides.
• **Supplementary Figure S3.** Examples of microbiome isolates lacking critical PT synthesis genes.
• **Supplementary Figure S4.** Workflow for PT-seq.
• **Supplementary Figure S5.** The frequencies of known PT- consensus sequences were low at boundaries of coding regions.
• **Supplementary Figure S6:** The density of PT-modified consensus sequences in CDSs and the boundaries of CDSs.
• **Supplementary Figure S7.** The distribution of PTs in *E. coli* BW25113.

• **Supplementary Table S1:** Gene neighborhood analysis of DndC protein. *Separate spreadsheet.*
• **Supplementary Table S2:** Gene neighborhood analysis of SspD protein. *Separate spreadsheet.*
• **Supplementary Table S3:** Dnd, Ssp, and Brx PT systems in 6616 BV-BRC bacterial genomes. *Separate spreadsheet.*
• **Supplementary Table S4:** PT dinucleotide analyses in human gut microbiome isolates using LC-MS*. Separate spreadsheet.*
• **Supplementary Table S5:** Dnd, Ssp, and Brx PT genes and other defense system genes in genomes of human gut micriobiome isolates. *Separate spreadsheet.*
• **Supplementary Table S6.** PT-seq read pileups in *B. salyersiae*. *Separate spreadsheet.*
• **Supplementary Table S7.** PT-seq read pileups in *B. faecalis*. *Separate spreadsheet.*
• **Supplementary Table S8.** PT-seq read pileups in *Lachnospiraceae sp. Separate spreadsheet.*
• **Supplementary Table S9**. Comparison of PT locations among classes of intergenic region, tRNA, rRNA and coding sequences (CDS). *Separate spreadsheet.*
• **Supplementary Table S10.** Bacterial strains used in the present studies.
• **Supplementary Table S11.** Primers used in the present studies.
• **Supplementary Table S12**: Query protein sequences used in this study. *Separate spreadsheet.*
• **Supplementary Table S13.** Mass spectrometry detection parameters for PT dinucleotides.
• **Supplementary Table S14.** PT dinucleotide high-resolution MS detection parameters used in this study.

### *Supplementary Figures*

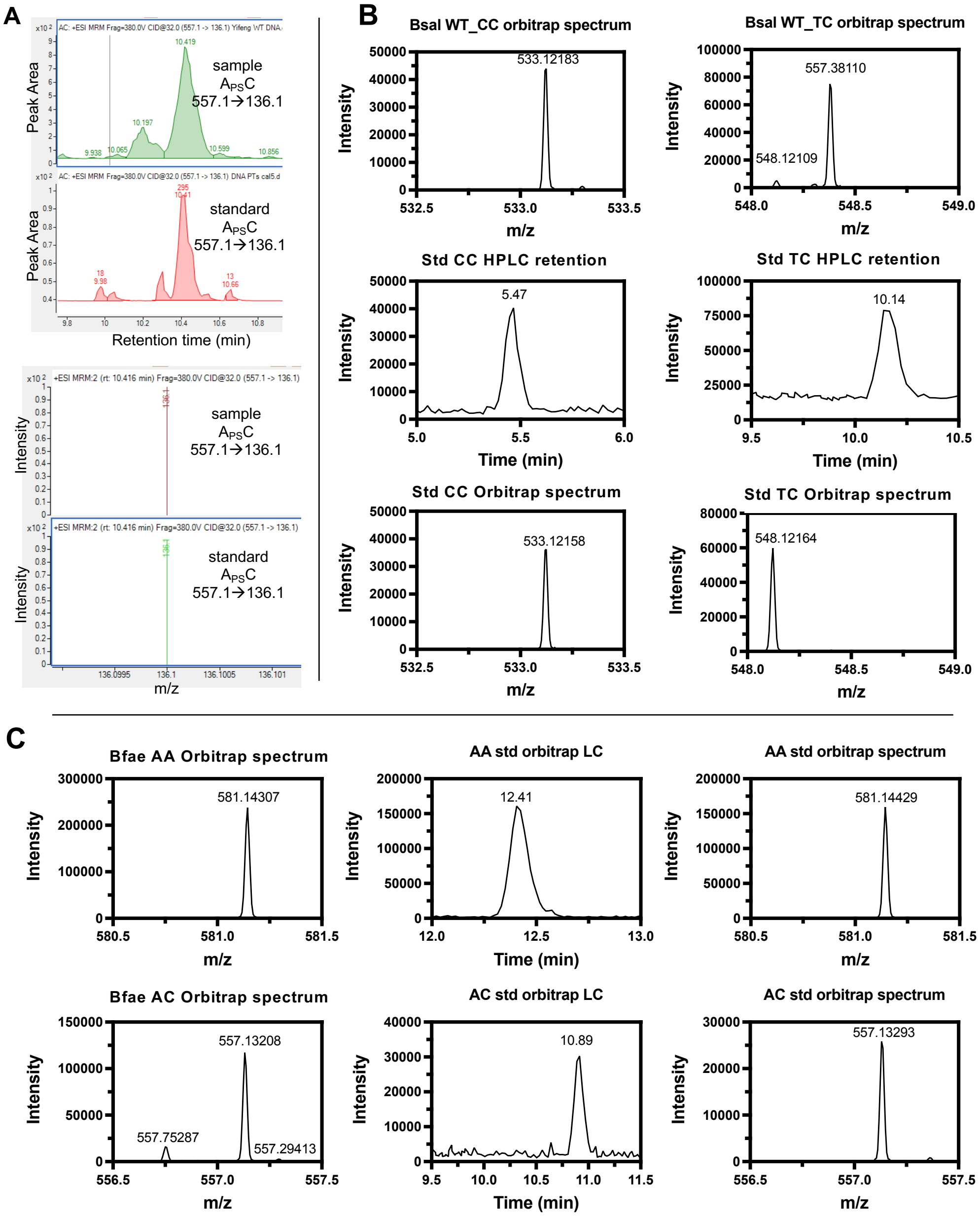


**Supplementary Figure S1**. Mass spectrometric validation of PT dinucleotides. (**A**) Comparison of the elution times and low-resolution mass transitions of $A_{PS}C$ PT dinucleotide in the *B. salyersaie* microbiome isolate ("sample") and synthetic standard ("standard"). (**B**) Orbitrap-derived exact masses of $C_{PS}C$ and $T_{PS}C$ PT dinucleotides in *B. salyersaie* microbiome isolate ("Bsal WT"") and HPLC retention and exact masses for synthetic standards ("Std"). (**C**) Orbitrap-derived exact masses of $A_{PS}A$ and $A_{PS}C$ in putative *Butyricimonas faecalis* isolate ("Bfae") and the exact masses and HPLC retention times for synthetic standards ("std").

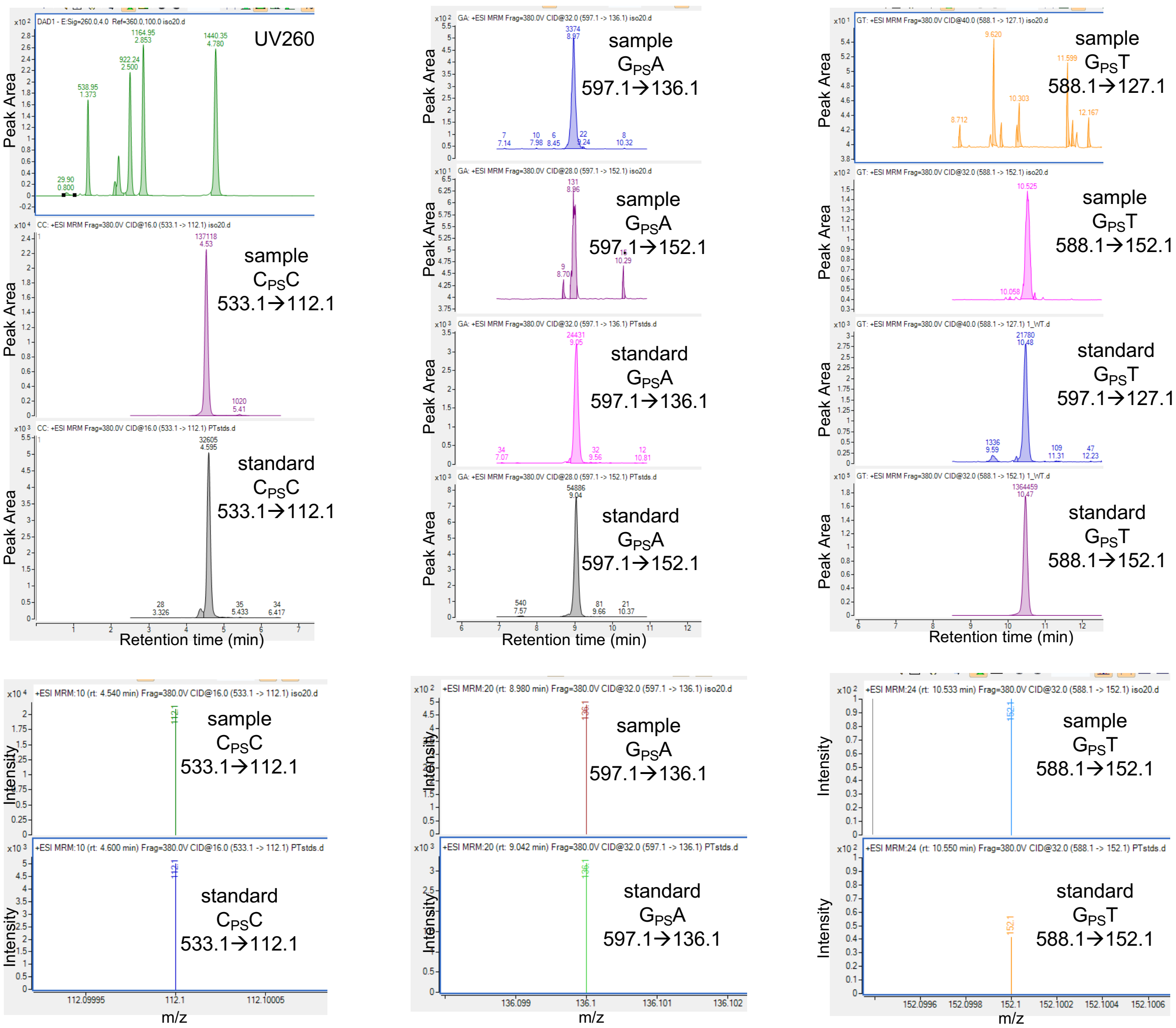


**Supplementary Figure S2**. LC-MS/MS characterization of PT dinucleotides. The elution time and mass of C$_{PS}$C (left), G$_{PS}$A (middle) and G$_{PS}$T (right) found in DNA from a *Prevotella* strain ("sample"; **Table S4**, Genome ID: 5645FP_0918_058_A7) and PT standards ("standard").

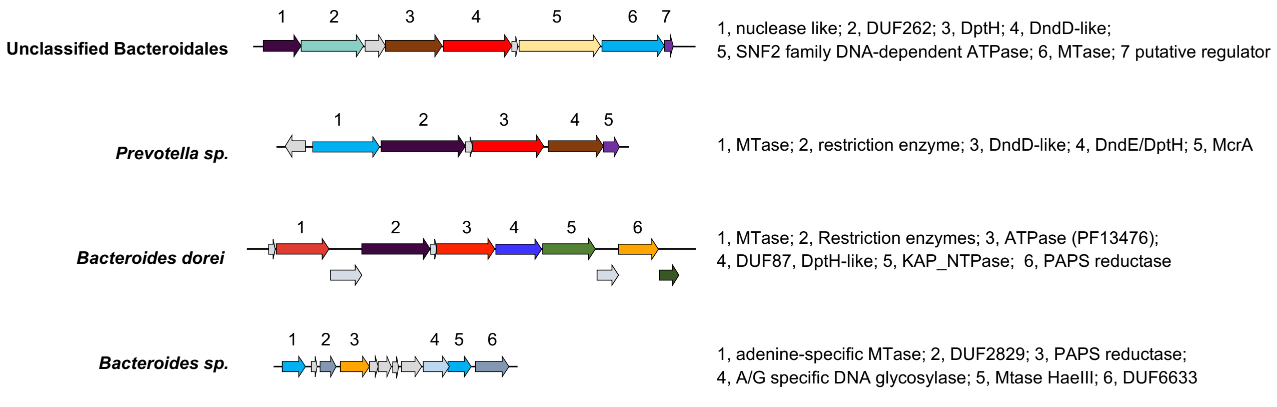


**Figure S3.** Examples of microbiome isolates lacking critical PT synthesis genes. The four examples shown lack one or more of the essential enzymes needed for PT synthesis: *dndCD*, *sspBCD,* and *brxPC* genes.

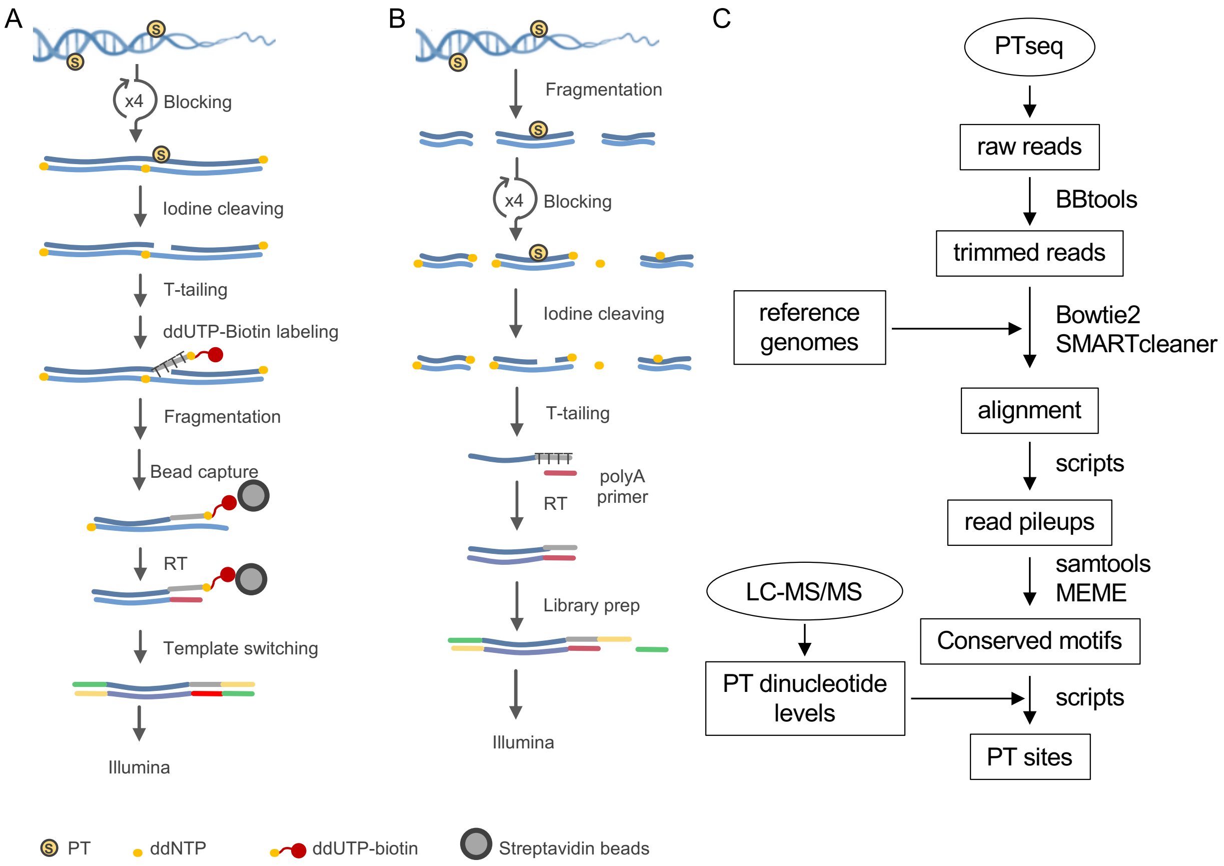


**Figure S4**. Workflow for PT-seq. (**A**) An upgraded PT-seq protocol. Pre-existing 3'-ends are blocked by four cycles of denaturation and blocking using terminal transferase and ddNTP. After iodine cleavage at PT sites, newly introduced 3'-ends are labeled with polyT tails and biotin-ddUTP. After sonication to fragment the DNA, biotinylated fragments are captured on streptavidin beads followed by library preparation using template switching and PCR technologies. (**B**) An earlier PT-seq protocol without biotin-labeling and streptavidin bead capture. (**C**) The workflow of PT-seq data analysis.

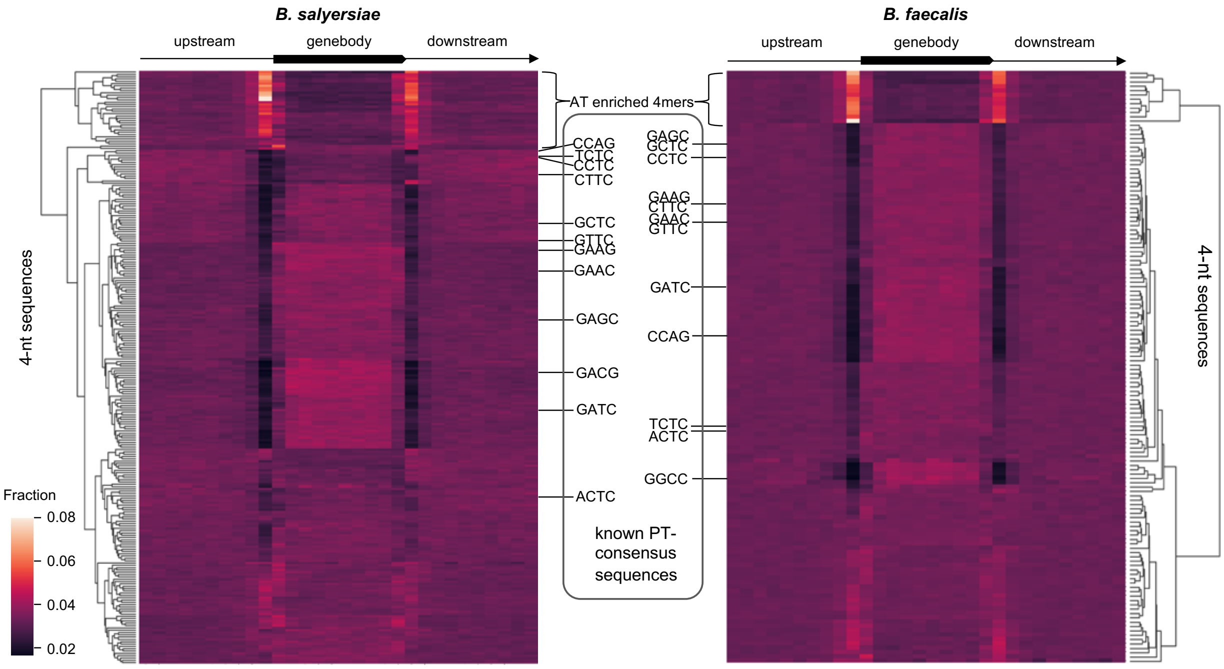


**Figure S5.** The frequencies of known PT- consensus sequences were low at boundaries of coding regions. The heat map of the distribution of 4- nucleotide sequence in all CDSs and their 1kb upstream and downstream regions in *B. salyersiae* (**left**) and *B. faecalis* (**right**). Each row represents the distribution of each of 256 4-nucleotide sequences from AAAA to TTTT in CDSs and their 1kb upstream and downstream regions. The distribution was colored according to the frequency of each 4-nucleotide sequence in each position. The known PT- consensus sequences, such as GAAC/GTTC in *E.coli*, GAGC/GCTC, and ACTC/CCTC/TCTC identified in this study are indicated by lines. The density and clustering were plotted using the heatmap package in R.

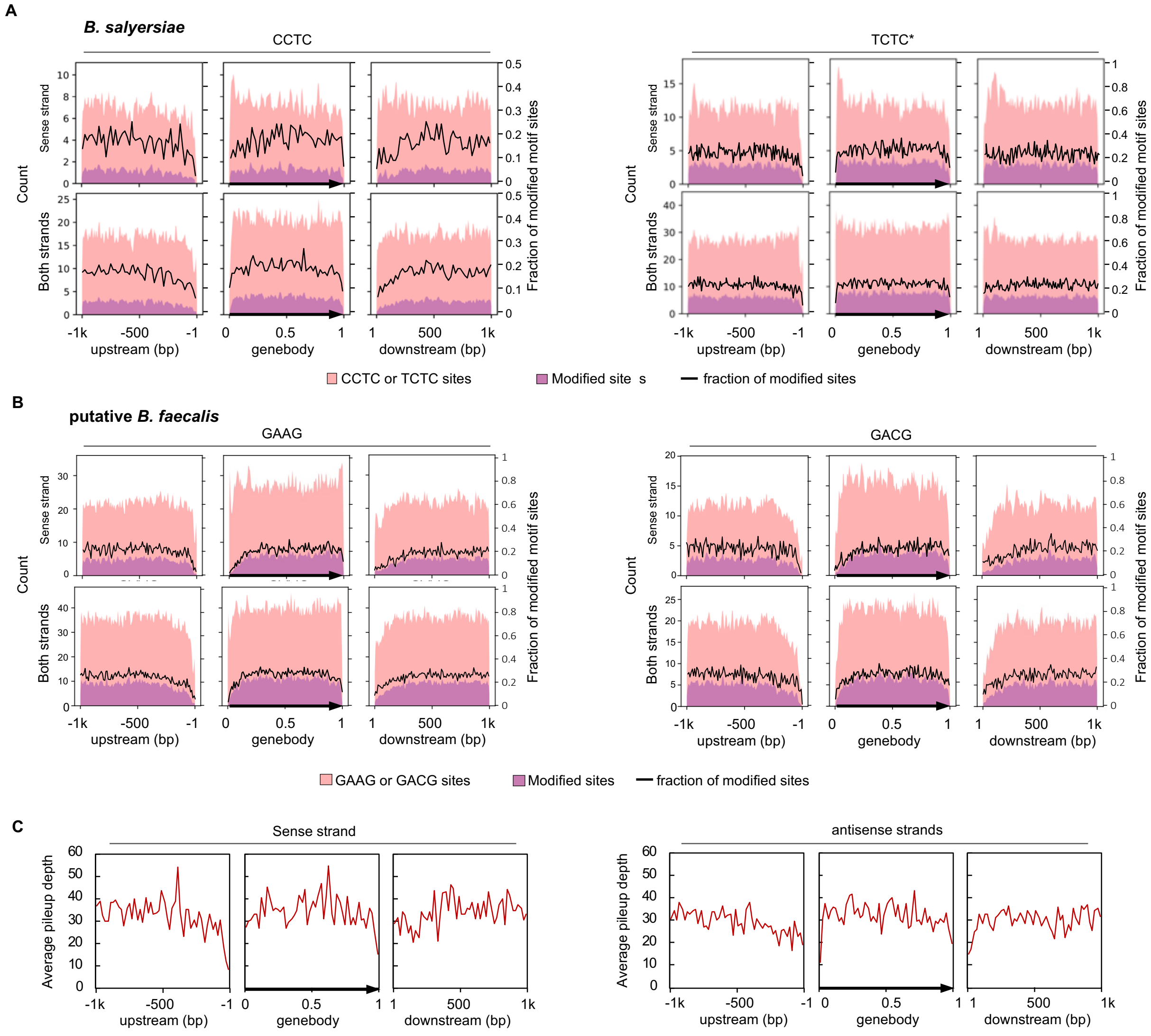


**Figure S6.** The density of PT-modified consensus sequences in CDSs and the boundaries of CDSs. The number of total consensus sequences in CDSs and their 1kb upstream and downstream regions in *B. salyersiae* (**A**) and putative *B. faecalis* (**B**), respectively*.* The number of consensus sequences in the sense strand (upper row) or both strands (lower row) are indicated in pink. The numbers of PT-modified consensus sequences are indicated in purple. The fraction of PT-modified sites is indicated by the black line. The asterisk denotes that a less stringent cutoff of depth (50) was applied for better sensitivity to plot a smooth curve for $T_{PS}CTC$ sites. (**C**) The average depth of read pileups that start with at least 50 reads in *B. salyersiae.* There is no significant difference between sense strand and antisense strand.

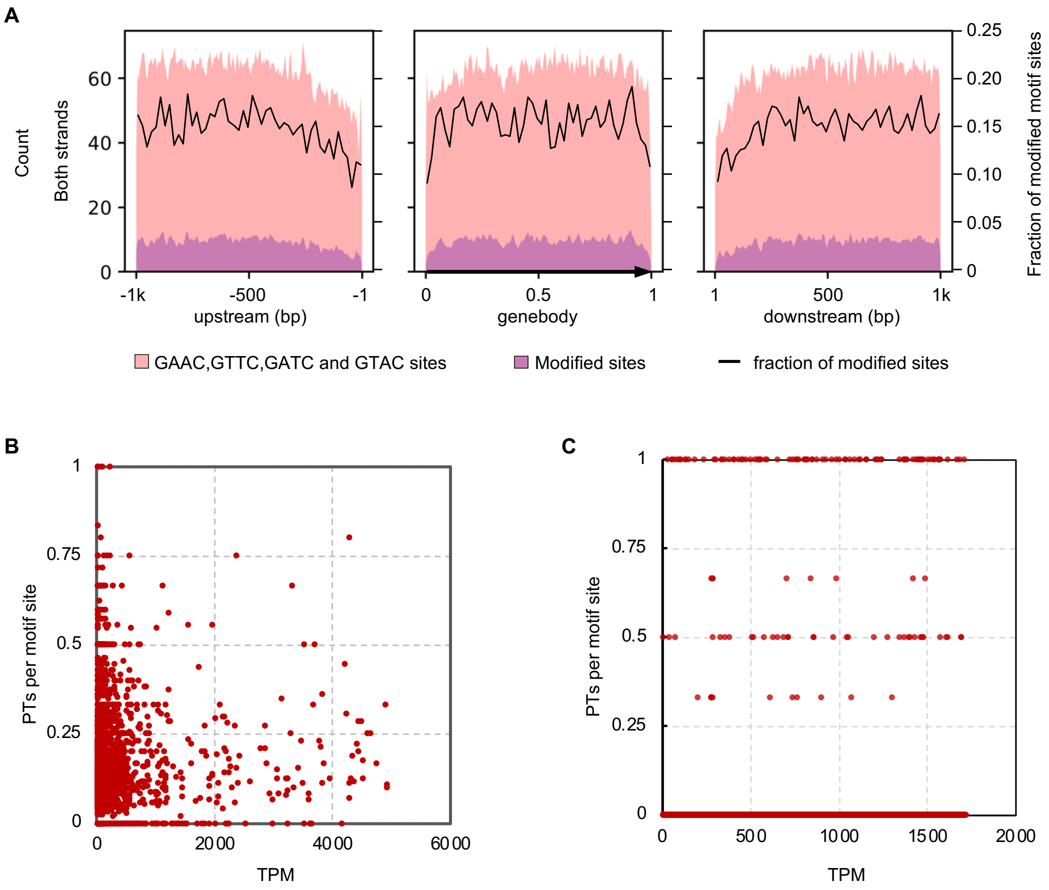


**Figure S7.** The distribution of PTs in *E. coli* BW25113. (**A**) The number of GAAC, GTTC, GATC and GTAC consensus sequences in both strands of CDSs and their 1kb upstream and downstream regions in *E. coli* BW25113. The number of total GAAC, GTTC, GATC and GTAC sites are indicated in pink. The PT-modified sites are plotted in purple. The fraction of PT-modified motif sites is indicated by the black line. (**B**) The plot of TPM (Transcripts Per Million) and PTs per GAAC, GTTC, GATC and GTAC motifs present in transcripts in *E. coli* BW25113. Data shown here are for 3227 transcripts that contain at least one motif site, 10 reads of RNA-seq data, at least 5 TPM, and less than 5000 TPM. (**C**) A plot of TPM and PTs per GAAC, GTTC, GATC and GTAC motifs within 50 bp upstream of start codons in transcripts in *E. coli* BW25113. Transcripts that contain at least one motif site within 50 bp upstream of start codons are shown.

### *Supplementary Tables*

**Supplementary Data Table S10.** Bacterial strains and plasmids used in this study.

| **Strain** | **source** |
|---|---|
| *B. salyersaie* DSM 18765 | DSMZ repository |
| *B. salyersaie* DSM 18765 Δ*mcrA* (HMPREF1532_02792) | this study |
| *B. salyersaie* DSM 18765 Δ*brxC* (HMPREF1532_02793) | this study |
| *B. salyersaie* DSM 18765 Δ*brxC* with pKF48 | this study |
| *B. salyersaie* DSM 18765 Δ*brxC* Iwith pKF49 | this study |
| *B. salyersaie* DSM 18765 Δ*brxC* with pKF58 | this study |
| *B. salyersaie* DSM 18765 Δ*brxC* with pKF59 | this study |
| *B. salyersaie* DSM 18765 Δ*brxZ* (HMPREF1532_02794) | this study |
| *B. salyersaie* DSM 18765 Δ*brxL* (HMPREF1532_02795) | this study |
| *B. salyersaie* DSM 18765 ΔHMPREF1532_02796 | this study |
| *Bacteroides thetaiotaomicron* VPI 5482 | Zitomersky *et al*.[1] |
| *Bacteroides thetaiotaomicron* VPI 5482 with pKF48 | this study |
| *Bacteroides thetaiotaomicron* VPI 5482 with pKF49 | this study |
| **Plasmid** | **source** |
| pLGB13 | Garcia-Bayona *et a*l.[2] |
| pFD340 | Smith *et al*.[3] |
| pKF58 (DPCBNLOC_02819 in pFD340) | this study |
| pKF59 (HMPREF1070_00305 in pFD340) | this study |
| pKF49 (HMPREF1532_02791 - 02793 in pFD340) | this study |
| pKF48 (HMPREF1532_02793 in pFD340) | this study |


[1]Zitomersky NL, Coyne MJ, Comstock LE. Longitudinal analysis of the prevalence, maintenance, and IgA response to species of the order Bacteroidales in the human gut. Infect Immun. 2011 May;79(5):2012-20. doi: 10.1128/IAI.01348-10.

[2]García-Bayona L, Comstock LE. Streamlined Genetic Manipulation of Diverse *Bacteroides* and *Parabacteroides* Isolates from the Human Gut Microbiota. mBio. 2019 10:10.1128/mbio.01762-19.

[3]Smith, C. J., Parker, A. Rogers, M. B. Plasmid transformation of *Bacteroide* spp. by electroporation. *Plasmid.* 1990 24, 100–109.

**Supplementary Data Table S11.** Primers used in the present studies.

| Purpose | Direction | Sequence |
|---|---|---|
| Construction of *B. salyersaie* DSM 18765 Δ*mcrA* (HMPREF1532_02792) | left flank forward | taagattagcattatgagtggggttgagacatcgggtattaag |
| | left flank reverse | ttgaaattcagttcattacgctgtcaatattttaag |
| | right flank forward | cgtaatgaactgaatttcaattttatacttgacaatgg |
| | right flank reverse | cgaattcctgcagcccggggggctgacggatgagaaatc |
| Comstruction of *B. salyersaie* DSM Δ*brxC* (HMPREF1532_02793) | left flank forward | taagattagcattatgagtgggcttggactaaagacgc |
| | left flank reverse | atgattcgagcgttgccattgtcaagtataaaattg |
| | right flank forward | aatggcaacgctcgaatcatcaaataaatagaag |
| | right flank reverse | cgaattcctgcagcccggggctgtttcgcaatcaaacc |
| Comstruction of *B. salyersaie* DSM Δ*brxZ* (HMPREF1532_02794) | left flank forward | taagattagcattatgagtgccaccatacgggctgtcg |
| | left flank reverse | tagcacccgtgttgaacgactgatacataggctac |
| | right flank forward | gtcgttcaacacgggtgctaccgaagatg |
| | right flank reverse | cgaattcctgcagcccggggctgactcacgaaagttatgagg |
| Comstruction of *B. salyersaie* DSM Δ*brx* (HMPREF1532_02795) | left flank forward | taagattagcattatgagtgcgctaattctgcatctttc |
| | left flank reverse | cttccataccacgaatcttatctgataagg |
| | right flank forward | taagattcgtggtatggaagaaaatgagttattttaagg |
| | right flank reverse | cgaattcctgcagcccgggggcggtcctgttccagtaatg |
| Comstruction of *B. salyersaie* DSM ΔHMPREF1532_02796 | left flank forward | taagattagcattatgagtgccttataggtcagttgatg |
| | left flank reverse | attacccccaaagcttatccaaattttccatatg |
| | right flank forward | ggataagctttgggggtaatttaacttgctggtaac |
| | right flank reverse | cgaattcctgcagcccggggagcgtccaccggtaccattc |
| Clone HMPREF1532_02793 into pFD340 | forward | aatcagaattgactctagaggaatttcaattttatacttgacaatg |
| | reverse | attcgagctcggtacccgggctatttatttgatgattcgagaattttatc |
| Clone HMPREF1532_02791-02793 into pFD340 | forward | aatcagaattgactctagaggcagaactgctcaatgttg |
| | reverse | attcgagctcggtacccgggctatttatttgatgattcgagaattttatc |

**Supplementary Data Table S13.** Mass spectrometry detection parameters for PT dinucleotides.

| Nucleosides | Precursor Ion | Product Ion | Retention Time (min) | Collision Energy | Cell Accelerator Voltage |
|---|---|---|---|---|---|
| d(ApsA) | 581.1 | 136.1 | 12.53 | 40 | 3 |
| d(ApsC) | 557.1 | 136.1 | 11.01 | 32 | 3 |
| | 557.1 | 112.1 | 11.01 | 24 | 3 |
| d(ApsG) | 597.1 | 152.1 | 9.7 | 48 | 5 |
| | 597.1 | 136.1 | 9.7 | 36 | 1 |
| d(ApsT) | 572.1 | 136.1 | 14.75 | 32 | 1 |
| | 572.1 | 127.1 | 14.75 | 48 | 5 |
| d(CpsA) | 557.1 | 136.1 | 8.29 | 32 | 1 |
| | 557.1 | 112.1 | 8.29 | 28 | 3 |
| d(CpsC) | 533.1 | 112.1 | 4.6 | 16 | 5 |
| d(CpsG) | 573.1 | 152.1 | 5.11 | 32 | 1 |
| | 573.1 | 112.1 | 5.11 | 32 | 5 |
| d(CpsT) | 548.1 | 127.1 | 8.79 | 36 | 5 |
| | 548.1 | 112.1 | 8.79 | 28 | 5 |
| d(GpsA) | 597.1 | 152.1 | 9.1 | 28 | 3 |
| | 597.1 | 136.1 | 9.1 | 32 | 1 |
| d(GpsC) | 573.1 | 152.1 | 7.64 | 32 | 5 |
| | 573.1 | 112.1 | 7.64 | 40 | 3 |
| d(GpsG) | 613.1 | 152.1 | 7.14 | 36 | 3 |
| d(GpsT) | 588.1 | 152.1 | 10.64 | 32 | 5 |
| | 588.1 | 127.1 | 10.64 | 40 | 3 |
| d(TpsA) | 572.1 | 136.1 | 12.92 | 28 | 3 |
| | 572.1 | 127.1 | 12.92 | 44 | 3 |
| d(TpsC) | 548.1 | 127.1 | 10.19 | 44 | 1 |
| | 548.1 | 112.1 | 10.19 | 24 | 5 |
| d(TpsG) | 588.1 | 152.1 | 9.2 | 32 | 5 |
| | 588.1 | 127.1 | 9.2 | 48 | 1 |
| d(TpsT) | 563.1 | 127.1 | 14.43 | 40 | 5 |
| dA | 252.1 | 136.1 | 5.23 | 12 | 1 |
| dC | 228.1 | 112.1 | 1.45 | 16 | 1 |
| dG | 268.1 | 152.1 | 2.71 | 4 | 3 |
| dT | 243.1 | 127.1 | 3.22 | 12 | 1 |

**Supplementary Data Table S14.** PT dinucleotide high resolution detection parameters.

| Nucleosides | Calculated Exact Mass | Retention Time (min) |
|---|---|---|
| d(ApsA) | 581.14388 | 12.41 |
| d(ApsC) | 557.13264 | 10.92 |
| d(ApsG) | 597.13879 | 9.61 |
| d(ApsT) | 573.13231 | 14.77 |
| d(CpsA) | 557.13264 | 8.27 |
| d(CpsC) | 533.12141 | 5.47 |
| d(CpsG) | 573.12756 | 5.31 |
| d(CpsT) | 548.12108 | 8.81 |
| d(GpsA) | 597.13879 | 9.61 |
| d(GpsC) | 573.12756 | 7.65 |
| d(GpsG) | 613.13371 | 7.2 |
| d(GpsT) | 588.12722 | 11.54 |
| d(TpsA) | 573.13231 | 12.88 |
| d(TpsC) | 548.12108 | 10.14 |
| d(TpsG) | 588.12722 | 9.18 |
| d(TpsT) | 563.12074 | 14.43 |
| dA | 252.10912 | 5.23 |
| dC | 228.09788 | 1.45 |
| dG | 268.10403 | 2.71 |
| dT | 243.09755 | 3.22 |
